# A quantitative analysis of the interplay of environment, neighborhood and cell state in 3D spheroids

**DOI:** 10.1101/2020.07.24.219659

**Authors:** Vito RT Zanotelli, Matthias Leutenegger, Xiao-Kang Lun, Fanny Georgi, Natalie de Souza, Bernd Bodenmiller

## Abstract

Cells react to their microenvironment by integrating external stimuli into phenotypic decisions via an intracellular signaling network. Even cells with deregulated signaling can adapt to their environment. To analyze the interplay of environment, neighborhood, and cell state on phenotypic variability, we developed an experimental approach that enables multiplexed mass cytometric imaging to analyze up to 240 pooled spheroid microtissues. This system allowed us to quantify the contributions of environment, neighborhood, and intracellular state to phenotypic variability in spheroid cells. A linear model explained on average more than half of the variability of 34 markers across four cell lines and six growth conditions. We found that the contributions of cell-intrinsic and environmental factors are hierarchically interdependent. By overexpression of 51 signaling protein constructs in subsets of cells, we identified proteins that have cell-intrinsic and extrinsic effects, exemplifying how cell states depend on the cellular neighborhood in spheroid culture. Our study deconvolves factors influencing cellular phenotype in a 3D tissue and provides a scalable experimental system, analytical principles, and rich multiplexed imaging datasets for future studies.

## 2 Introduction

The ability of a cell to sense and adapt to its local environment depends on an intracellular signaling network that integrates paracrine, juxtacrine, nutritional, and mechanical cues among others to drive phenotypic decisions (Fig. 1a). Genomic alterations that deregulate environment sensing and signaling can enable cells to grow outside their physiologically permissive tissue context, leading to diseases such as cancer. However, analyses of cells grown in two-dimensional culture indicate that even strongly deregulated cells depend on and react to microenvironmental cues [1, 2]. Variability introduced by microenvironment-induced cellular plasticity may contribute to the clinically relevant tumor cell heterogeneity observed in cancer tissues [3, 4].

**Figure 1.**
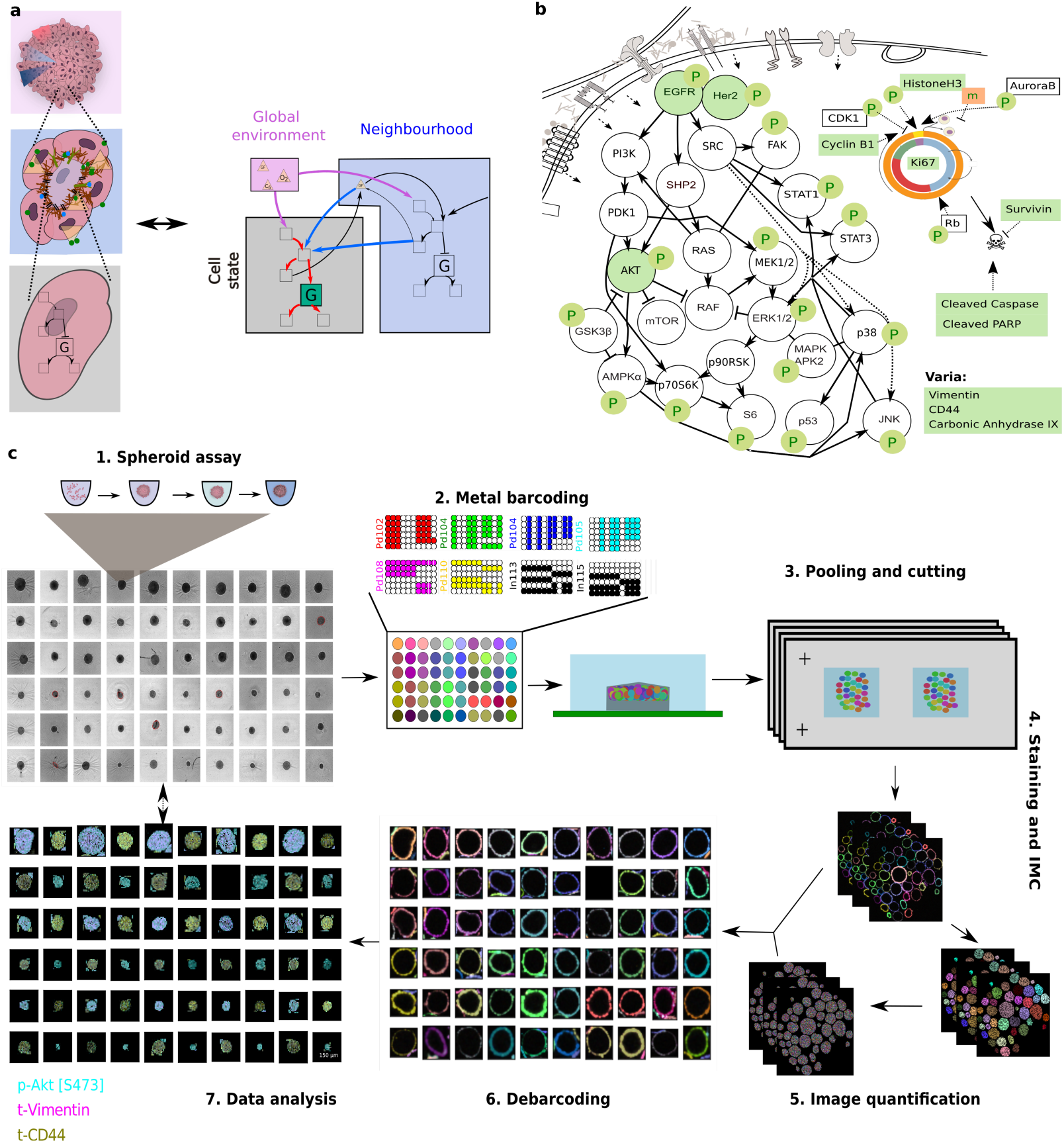
Barcoded IMC assays allow efficient spatial profiling of pooled spheroids. **a** Cells sense their environment and compute cellular decisions via a signaling network. The schematics (left) depict spheroids at different scales: spheroid with global gradients of, for example, nutrients and oxygen (top), cellular neighborhood (middle), and single cell (bottom). The models (right) depict the interdependence of global environment (pink box), local neighborhood (blue box), and intracellular state (grey box) in determining levels of a given marker, G. **b** A schematic illustration of the signaling network markers, cell-state markers, and other phenotypic markers measured using IMC (green). Nodes depicted in white were not measured. **c** Flow diagram of barcoding approach used for multiplexed IMC analyses of spheroids.

Assessments of spatial tumor cell heterogeneity based on protein and transcript measurements have been generated for several types of tumors [5–10]. Missing, however, is a quantitative understanding of how the three-dimensional (3D) tissue environment influences heterogeneity. Existing atlases of cancer tissues are based on static measurements of cellular markers that cannot reliably discriminate environment-dependent phenotypic plasticity from phenotypic variation due to genomic or lineage differences [5, 9, 11]. To quantify variability caused by the environment, it is necessary to identify comparable cells that vary phenotypically only because their environments differ.

To generate a dataset consisting of such comparable cells, we cultured and analyzed spheroids consisting of clonal cells. We used an established method to grow cell lines as multicellular spheroids [12]. We reasoned that this type of system would allow quantification of the influence of the global environment, local environment, and cell state on measurable cellular phenotypes. Further, as spheroid cell culture is compatible with 96-well microplates, this technology is suitable for large-scale perturbation studies and can be extended to more complex co-culture systems or heterocellular organoids [13–16].

To efficiently quantify phenotypic and signaling states of cells in spheroids at high-throughput, we coupled metal-based barcoding with antibody-based multiplexed imaging mass cytometry (IMC) [17–19]. This approach allowed us to process up to 240 spheroids simultaneously and to measure the levels of dozens of phenotypic markers in hundreds of sphere slices containing hundreds of thousands of cell sections. We evaluated spheroids formed by four cell lines, each grown in six different growth conditions, quantified single-cell variability, and analyzed how cell state, local neighborhood, and global environment interact to contribute to cell-to-cell variability in marker expression. Further, to explicitly probe cell-to-cell signaling interactions, we developed a chimeric overexpression-based approach to test the effects of overexpression of 51 ligand and receptor components of more than a dozen different signaling pathways on responses of neighboring cells. Our approach provides a blueprint for large-scale studies on any type of 3D microtissues and for deconvoluting microenvironmental and internal contributions to cellular phenotype in spatial data.

## 3 Results

### 3.1 Spheroid culture coupled with multiplexed imaging enables quantification of phenotypic variability

To investigate factors that influence phenotypic variability in spheroids consisting of clonal cells, we developed a combined experimental and computational workflow. We grew cells as spheroids and imaged histological sections of these 3D tissues using IMC [19] with a panel of antibodies that detect 20 growth-signaling markers, nine cell-cycle or apoptosis markers, and four markers indicating molecular phenotype (Fig. 1b, Tab. S1). We characterized the internal state of each cell by quantifying marker levels in individual cell sections (Fig. 1a, intracellular network, grey box). We evaluated the local environment of a cell by quantifying marker levels within neighborhood cells (Fig. 1a, blue box). Finally, since this culture system shows radially symmetric gradients of nutrients, oxygen, and growth factors [12, 20, 21], we used the distance from a cell to the border of the spheroid as a surrogate measurement of global environmental influences on phenotype (Fig. 1a, violet box).

Histological sectioning, staining, and analysis of individual 3D microtissues is challenging to perform at scale: Cutting and staining spheres individually is very labor and resource intensive. To improve scalability, we adapted a metal-based barcoding approach from single-cell mass cytometry [22, 23] (Fig. 1c). This approach enabled barcoding of up to 240 single spheroids grown in individual wells of multi-well plates. After barcoding, spheres were pooled into a dense cylinder for efficient embedding and cutting. Sections from the spheroid plug were then imaged using brightfield imaging and sections containing dozens of spheres were selected for staining and IMC analysis. The metal barcodes allowed us to relate each imaged sphere section to its sphere of origin and thus to the cell line and perturbation. Pooled processing of spheres reduced the manual labor and staining of spatially concentrated spheres reduced the amount of antibody required compared to other approaches [24]. Finally, we improved data quality by applying rigorous quality control steps on the cell, sphere slice, and intact sphere by leveraging orthogonal imaging modalities such as brightfield and fluorescent imaging (Methods).

We grew spheroids from four widely used epithelial cell lines that reproducibly form smooth spheroids [25]. T47D cells are derived from a breast cancer tumor [26], HT29 and DLD1 lines are derived from colorectal tumors [27, 28], and T-REx-293 cells are derived from human embryonic kidney cells [29]. We chose these cell lines with the goal of identifying cell line-specific and general factors that influence phenotypic variability. In addition, to examine whether our results were affected by spheroid size or growth time, we grew each of these four cell lines at three cell seeding concentrations (5 replicate wells each) and for two different time periods (72 h and 96 h). After cutting the pooled spheroid pellets, sections were stained with our antibody panel (Fig. 1b, Tab. S1) and imaged using IMC. After quality control and image processing, our data included 517 cuts from 100 spheres, corresponding to 228’740 cell sections with an average of 19’530 cell sections per cell line and growth condition (min=1’426, max=28’170, Tab. S2).

### 3.2 Marker levels show strong dependence on environment and are cell intrinsically and spatially correlated

We segmented the imaged spheroids into single-cell sections using a combination of machine learning and computer vision algorithms and quantified the average level of each measured marker for each single-cell. Dimensionality reduction analysis showed a near perfect separation into cells of the different cell lines as identified by debarcoding (Fig. S2a). We further confirmed that there were likley no misassignments during debarcoding with a clustering-based analysis (Fig. S2b-d). Visual inspection of spheroid images showed clear marker-specific spatial variation. Certain markers appeared in patches of cells, whereas levels of other markers were dependent on distance to the spheroid border (Fig. 2a, b). These results indicate that both the local environment and global effects influence marker expression.

**Figure 2.**
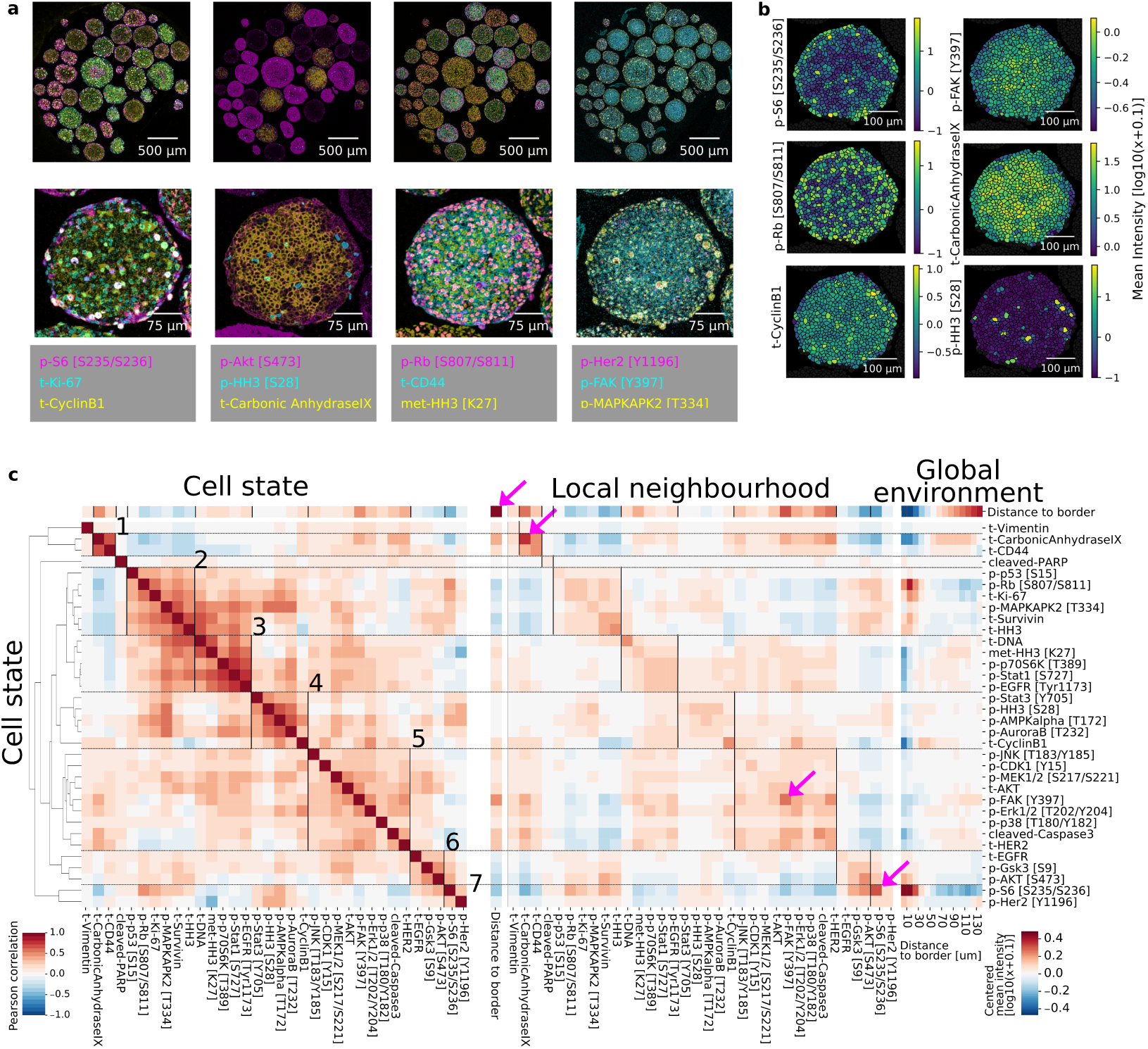
Multiplexed imaging captures spatial organization of spheroids. **a** Example IMC images of a spheroid plug (top row) and a HT29 spheroid section (bottom row). **b** Examples of image quantification showing log10-transformed average counts per cell section for the indicated markers. **c** Correlation analysis of HT29 spheroids (96 h growth). Left: Pearson correlation matrix of markers within each cell. Clusters (indicated by horizontal lines and labeled with numbers) are based on hierarchical clustering of the intracellular marker correlation (distance cosine, metric average linkage). Middle: Correlation matrix of markers in all cells (rows) and average marker levels in neighboring cells (columns). Right: Median log10 intracellular marker levels as a function of the distance to the spheroid border. Values centered around 0. Pink arrows highlight strong spatial autocorrelations (Pearson’s r > 0.5).

To systematically investigate intracellular, local, and global relationships for the 34 markers measured, we calculated Pearson correlations between intracellular levels of each marker in a given cell (cell state), as well as between intracellular markers and the average levels of markers in the immediate neighbors of the cell (local neighborhood). We also calculated the distance from the cell to the spheroid border as a proxy for the global environment and visualized average marker levels relative to this distance. The results for the HT29 cell line are representative (see Fig. S3a-c for data on all cell lines), and analyses of this cell line in one growth condition are discussed in this section.

Hierarchical clustering of the intracellular marker correlation matrix identified seven clusters (Fig. 2c, left). Clusters 2, 3, 5, 6 and 7 contain markers of activated growth signaling in the EGF and AKT/mTOR pathway, and cell-cycle markers. Mitotic markers are found in cluster 4. Cluster 1 consists of the classical hypoxia marker carbonic anhydrase 9 and the cell-cell adhesion marker CD44. Vimentin and cleaved PARP, which were expressed at very low levels in these spheres, did not cluster with other markers. Markers within the same cluster are consistently positive or negatively correlated with distance to border (Fig. 2c, top row), indicating that these intracellular states are linked to the spatial position within the sphere.

We next asked how these clusters mapped onto correlations between markers in neighboring cells. We correlated intracellular marker levels with the average marker levels of all cellular neighbors and ordered the resulting correlation heatmaps according to the clustering derived from intracellular marker correlations (Fig. 2c, middle). This ordering was in agreement with correlations between intracellular markers and average marker levels in neighboring cells, suggesting that intracellular marker correlations also capture correlations in the local neighborhood. Marker levels averaged over neighboring cells were even more strongly correlated to distance to spheroid border than were the intracellular levels, indicating that the local neighborhood is strongly dependent on the spatial position in the sphere.

We next focused on spatial autocorrelations (i.e., the correlations between an intracellular readout and the same readout in neighboring cells) (Fig. 2c, middle, entries on diagonal). Low autocorrelation is indicative of markers being locally variable, while high autocorrelation suggests either that a marker occurs in cell patches or varies smoothly in the local cell neighbourhoods. Most readouts had weak to medium spatial autocorrelation, but four had strong autocorrelations (pink arrows; Pearson’s r > 0.5). The strongest autocorrelation was found for the distance to border readout, our surrogate measurement for the global environment; unsurprisingly, this was almost perfectly correlated with the average distance-to-border of neighboring cells. The other three strongly auto-correlated markers, p-S6, carbonic anhydrase, and p-FAK, were also all highly correlated with the distance-to-border measure (Pearson’s r with distance to border > 0.5); these gradients of expression were confirmed visually in spheroid sections (Fig. 2a-b). Thus, spatial autocorrelation can capture effects of the global environment. However, low spatial autocorrelation of a marker does not necessarily imply a lack of influence by the global environment. For example, p-Rb, a marker of cells that have completed the G1/S transition, showed a strong distance-to-border effect (Fig. 2c, right), yet only a moderate autocorrelation (Pearson’s r= 0.35).

Direct visualization of average marker levels as a function of distance to border confirmed that clusters defined by intracellular correlations show similar marker localization patterns (Fig. 2c, right). This supports our hypothesis that intracellular marker correlations capture elements of the global environment (i.e., spatial position within the spheroid). We also observed a spatial segregation between markers of growth signaling, early cell-cycle, and late cell-cycle in all cell lines: AKT/mTOR signaling peaked in the outermost sphere layer, early cell-cycle markers (pRB, Ki67) were located in the penultimate layers, and markers of the late cell cycle (Cyclin B1) were more located in the middle layers of the sphere (Fig. S3d). These are additional examples of how cellular states are localized within the sphere environment and thus carry information about the spatial position of a cell. Taken together, our analysis indicates that intracellular markers are not only correlated to other markers that reflect cellular state, but that these states are also closely related to the cellular states of neighbors and the spatial location of cells in the global environment.

#### 3.2.1 Measurements of environment, neighborhood, and cell state are redundant

Given the strong and highly structured correlations observed, we asked to what degree marker levels are predictable by environment, local neighborhood, and cell state. We used linear modeling to predict the levels of each marker based on different predictive modules: the *global environment module* (a nonlinear function of the distance-to-border), the *local neighborhood module* (the average marker levels of direct neighbors without autocorrelation), the *local autocorrelation module* (average marker levels of the predicted marker in immediate neighbors), and the *internal cell state module* (all other internal markers) (Fig. 3a). In 56% of cases, the linear model including all modules explained more than 50% of the marker variability (Fig. 3b). With the exception of few highly cell line-specific markers, total marker variability explained was usually similar for the different cell lines. In the best cases, the model explained about 85% of the total variation. The residual unexplained variance likely reflects a combination of technical variability in staining, detection, and quantification, the biological variability, and the inability of the linear model to capture non-linear marker relationships. There was a clear relationship between average predictability and signal intensity for low-intensity markers (Fig. S4a), but not for markers expressed at medium to high intensity levels (higher than 1 average count per cell pixel). Thus, technical noise likely dominated the detection of the low-intensity markers.

**Figure 3.**
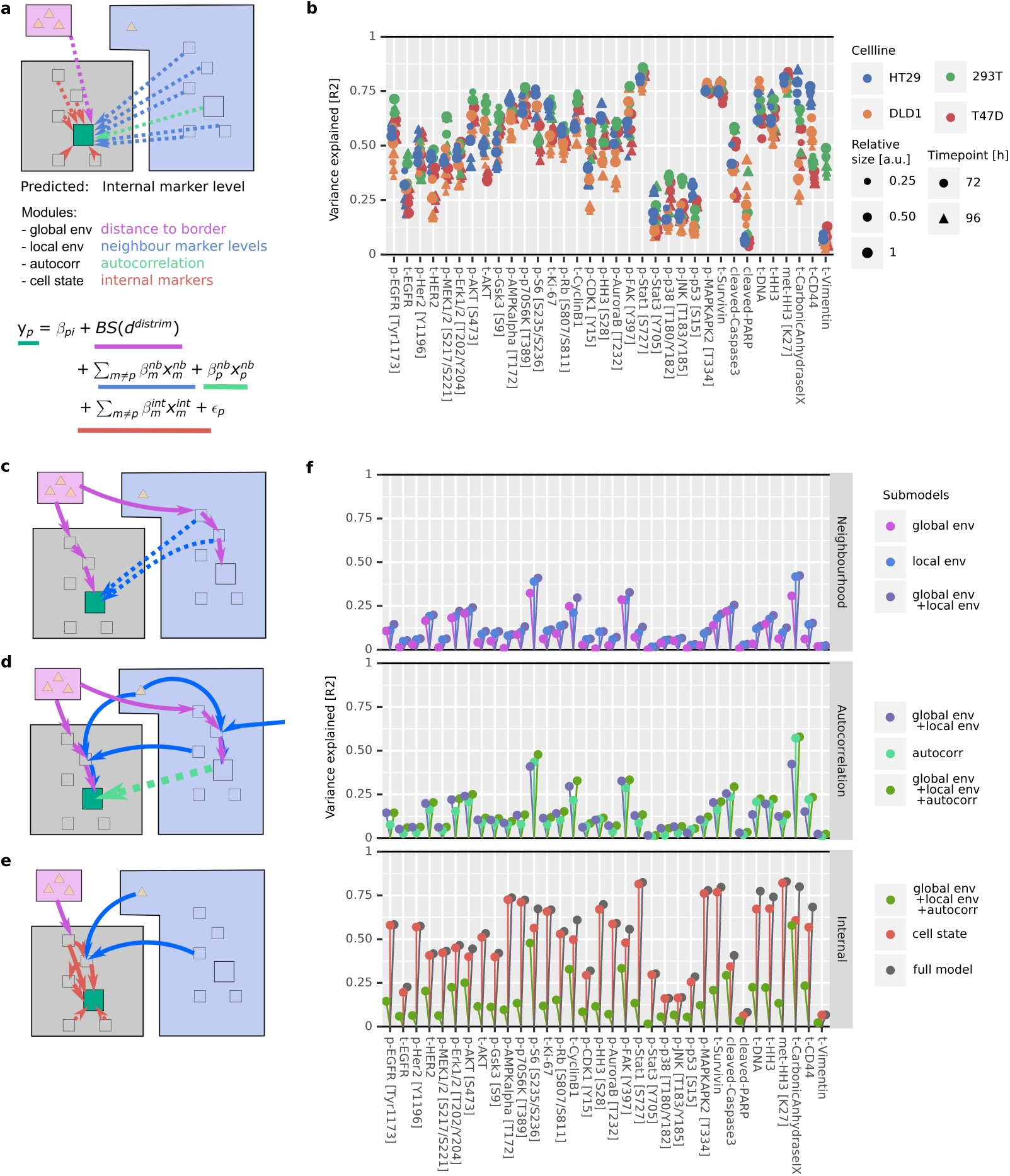
Global environment, local neighborhood, and cell-state are not independent predictors of single-cell marker levels in 3D spheroids. **a** Marker levels were predicted with a linear model using modules representing global environment (violet in schematic), local neighborhood (blue), autocorrelation (teal), and cell state (red). **b** Variance explained by the full model plotted for each marker, for all cell lines, and for all growth conditions. **c** The schematic depicts that markers in a given cell (green square) are directly correlated with neighboring cell markers due to mechanistic interactions (solid blue arrows) or indirectly correlated (dashed blue arrows) due to the global environment (violet arrows) affecting both cells and their neighbors. For all schematics (c - e), bold arrows indicate a direct effect and dotted arrows indicate indirect statistical correlations. **d** The schematic depicts that a marker (green square) strongly dependent on the local and global environment is statistically autocorrelated in neighboring cells (dashed teal arrow). **e** The schematic depicts that environmental influences affect marker levels via other intracellular proteins, so that certain internal markers capture environmental effects (red arrows). **f** Variance explained by the indicated modules for all markers in all cell lines and growth conditions. The data are visualized to illustrate the minimal added explanatory power of the local neighborhood over global environment (top), of autocorrelation over other spatial factors (middle), and of internal cell state markers over all environmental factors (bottom).

Next, we investigated the explanatory power of the individual (Fig. 3c-f). We expected that modules would not be independent in their explanatory power due to properties that result from the spatial tissue architecture: A cell and its neighbors, by virtue of their proximity, are subject to very similar global environmental cues. The global environment will thus similarly influence marker expression in a cell and its neighbors, leading to an indirect correlation between the two (Fig. 3c). We therefore expected that cell-state measurements of the local neighborhood should also capture marker variability caused by the global environment. This was strongly supported by our data: The linear model based on the global environment module alone explained a median of 7.6% of variation. The local neighborhood module alone explained a median of 12.8% of variation. Adding the global environment module to a model containing the local neighborhood module only improved the predictive power by a factor of 1.11 (+1.4%, Fig. 3f top, Fig. S4b). This indicates that indeed the local neighborhood largely captures the global environment in the ability to explain marker variation.

By similar reasoning, if expression of a marker in a given cell is strongly determined by the local and global environment, levels will be similar in neighboring cells (i.e., it is likely to be spatially autocorrelated). In this case, the local environment influences the expression of a given marker both in the cell of interest and in its neighbors (Fig. 3d), and autocorrelation alone should explain a substantial fraction of marker variation caused by local and global neighborhood effects. Supporting this hypothesis, local autocorrelation alone explained a median of 12% of marker variability in our data. The global environment and local neighborhood features together explained a median of 15% of marker variability. Adding these features to a model based on local autocorrelation improved the predictive power by 1.39 fold (+4.1%). This indicates that local autocorrelation alone captures more than half of the variability explained by spatial effects (Fig. 3f middle, Fig. S4b).

Finally, since cells convert external stimuli into an intracellular response via a highly interconnected intracellular signaling network, we expected that environmental effects should influence both expression of an environmentally determined marker and other related internal markers (Fig. 3e). Thus, a comprehensively measured internal cell state should capture much of the marker variability caused by the environment and neighborhood. This effect can be seen in our dataset: A model based solely on the internal cell state module captured more variability than a model with all neighborhood terms in almost all cases (97% of cases, Fig. 3f bottom, Fig. S4b). The internal cell state markers alone explained a median of 48% of variability, whereas all environmental terms together explained 17% of variability. Adding the environmental modules to the internal cell state module (to yield the full model) explained a median of only 1.05 fold more variability (+1.8%) than did the model based on the internal cell-state module alone.

In summary, a linear model based on measured global, local, and internal cell-state features predicted a substantial fraction of single-cell marker variance in homogenous 3D spheroids. Based on a conceptual model of cells interacting in tissue, we expected that the abilities of global environmental features, local environmental features, and intracellular features to predict marker variation would not be independent. We found that this expectation was strongly supported by our data. Furthermore, these interdependencies appeared to follow a hierarchy, where the explanatory power of the global environment is captured by that of the local environment, which in turn is captured by that of intracellular features.

#### 3.2.2 Step-wise regression identifies environmental marker dependencies

Next, we sought to compare the marker variation explained by global environment, local neighborhood, and internal cell state for each marker across cell lines and growth conditions. We reasoned that the additional variability explained as each module is added step-wise to a regression model would be biologically informative [30, 31]. The increasing order of explanatory power described above, which supports our conceptual model of cells interacting in tissue (Fig. 1a, Fig. 3c-e), suggests that sub-modules should be added in the order of global environment, local neighborhood, autocorrelation, and internal cell state (Fig. 4a). Exhaustive permutations confirmed that this sequence (of all possible sequences of step-wise addition of these modules) highlights the contributions of all factors optimally (Methods 7.3.1). Other sequences of module addition, by contrast, either mask the contributions of some modules or incorrectly exaggerate the contributions of others (Fig. S4c). In other words, with this sequence of addition, each added module approximately captures the variability explained by the modules added previously, again suggesting that we have identified interdependent predictors of marker variation that follow a hierarchical order of their explanatory power.

**Figure 4.**
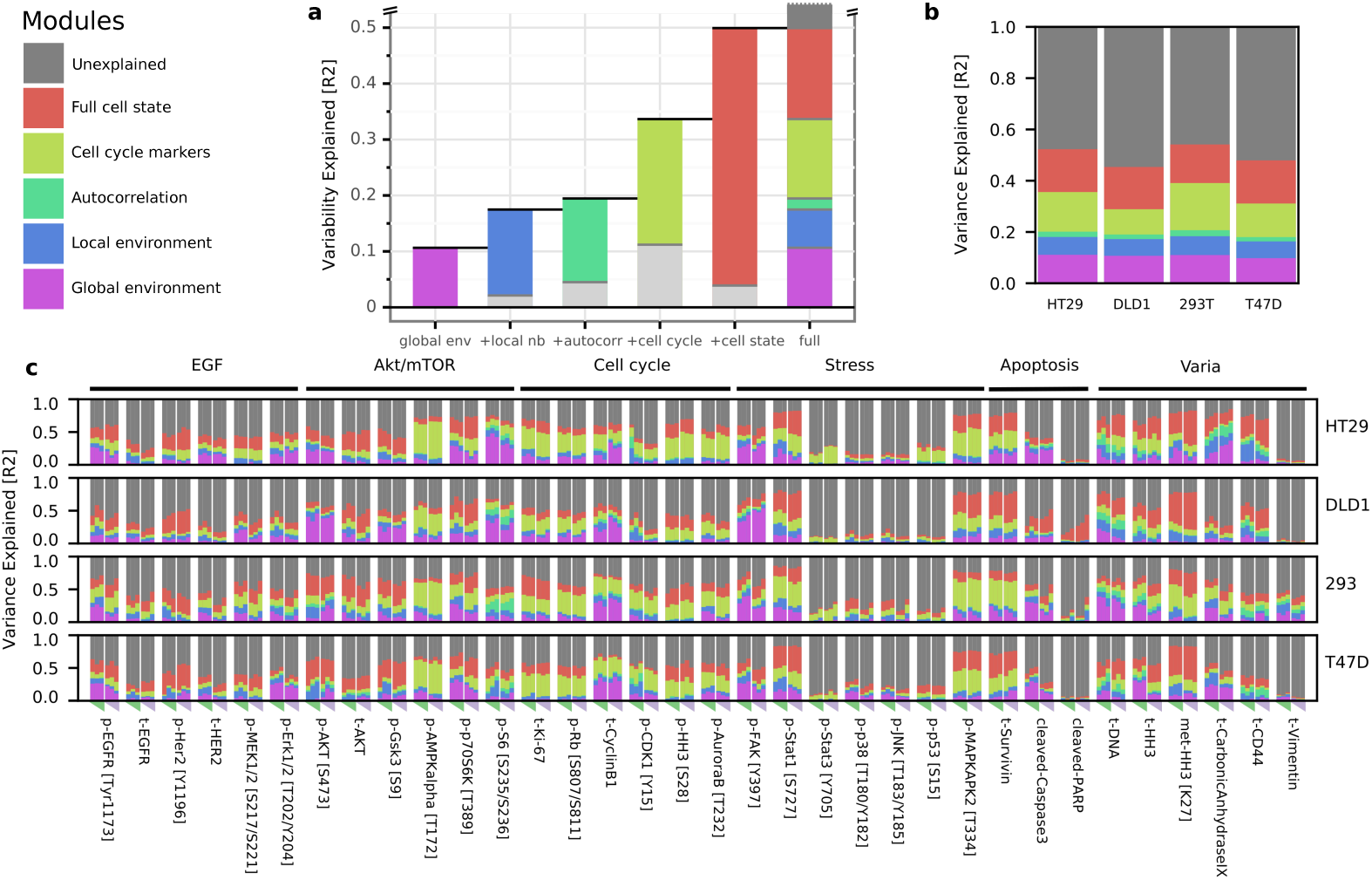
Marker variance is explained by cell-intrinsic and environmental factors. **a** Average marker variability over all markers explained by global environment, local neighborhood, autocorrelation, cell-cycle markers, and all intracellular markers. For each bar, colored portions indicate the variability explained by the particular module alone and the light gray portion indicates the additional variability explained when the previous module or modules are also included in the model. Since the variability explained by each feature is not additive but roughly follows a hierarchy, the contributions to the full model are represented as a stacked barplot. **b** Contributions of the different modules to marker variance for each cell line, averaged over all markers and growth conditions. **c** Contributions of the different modules to marker variance for each cell line and growth condition. Rows represent cell lines, columns show marker abundance at a specific growth condition. Columns represent growth conditions varied by sphere sizes (triangle, 0.25/0.5/1.0x cells) and growth time (72 h green, 96 h pink).

The cell cycle is a major source of cell-to-cell variability [32–34]. To specifically highlight variability linked to cell-cycle, we classified the internal markers into cell-cycle and non-cell-cycle markers (Tab. S1).

Averaged over all cell lines, markers and conditions, the linear model containing all modules explained 50% of variation, whereas 20% was explainable by all environmental factors (Fig. 4a). Within the spatial effects (i.e. global environment, local neighborhood, and local autocorrelation), the global environment explained on average 55% of variability. Averaging across all markers and growth conditions for each of the four cell lines showed similar dependencies (Fig. 4b), suggesting that each of these cell lines on average reacts similarly to internal and environmental influences when grown as 3D spheroids.

Our concise visualization based on the hierarchy of explanatory power also enabled fine-grained comparison of how each of the 34 markers depend on the global and local environments in four cell lines and under six growth conditions, allowing more than 4000 comparisons (Fig. 4c). We note that, across the dataset, the average standard deviation of the explained variability was less than 0.04 for all models (overall average 0.038, iqr. 0.02-0.049) across five spheroid replicates for each of the 24 growth conditions. We observed both general and cell line-specific effects.

Focusing on cell-cycle markers, we found that for a given marker, an average of around 50% of variability was explained by the full model. Cell cycle markers and environment together captured around 75% this variation. An exception across all cell lines was p-HH3, a mitotic marker, for which environment and cell-cycle markers explained only 58% of all explainable variation. This suggests that mitosis is a state particularly strongly linked to the cellular state as a whole and not only to cell-cycle markers. In another example of cell line-specific effects, Ki67 variability was strongly linked to non-cell-cycle intracellular markers specifically in T-REx-293 cells. Early cell-cycle markers phosphorylated RB and Ki67 showed little dependence on global environment in T47D cells (approximately 2% variability explained); in comparison, in HT29, DLD1, and T-REx-293 cells, the global environment explained 7-12% of the variability of these markers. Of all the cell-cycle markers, cyclin B1 levels were the most dependent on the environment, with an average of more than 20% of variability explainable by global environmental gradients in all cell lines.

The AKT/mTOR pathway is involved in growth and nutrient signaling [35]. We found that levels of multiple markers of this pathway are explained by environment and local neighborhood features. A downstream readout of this pathway, p-S6, was strongly dependent on environmental factors in all cell lines. Other upstream markers, such as p-AKT and p-GSK3beta, showed cell line-specific effects: Environmental factors had higher explanatory power for levels of these markers in DLD1 than other cell lines. The variability of total AKT was more linked to cell state than environment. Finally, levels of p-AMPK, reported to be a nutrient sensor [36], was strongly explained by the cell-cycle but only slightly by environmental factors. This is also reflected in the correlation maps, which showed that p-AMPK expression was correlated with that of mitosis markers (Fig. 2c, S3a-c), consistent with the reported association of this marker with the mitotic spindle [37, 38].

In summary, our analysis of multiplex imaging data in homogenous 3D tissue models allowed a detailed deconvolution of the factors affecting marker variation. Internal and environmental modules predict on average around 50% (up to 85%) of marker variability. The variability explained by different factors was on average similar across cell lines. There were, however, impacts of cell line and growth conditions on expression levels of certain markers. Our data allow granular identification of these cell line- and growth condition-specific patterns in marker dependencies.

### 3.3 Signaling deregulation affects cells and their neighbors in spheroids

Our experiments showed that, after correcting for global effects, on average 6% of marker variability was predicted by neighboring cell markers. To explore whether these correlations reflect spatial coordination due to active cell-to-cell communication, other biological effects, or technical artifacts, we developed an overexpression system to induce changes in individual cells and investigate the effects on neighboring cells. We hypothesized that in the overexpression context, active cell communication should lead to systematic changes in neighboring cell states. We used a previously described library of 32 pro-cancer-signaling protein constructs involved in 17 pathways and containing many common cancer driver mutations [39]. We supplemented this library with ten growth factor receptors, nine ligands, and four negative controls (Suptable. S3). Each clone was transformed into an inducible, GFP-tagged expression vector and individually transiently transfected into separate wells of T-REx-293 cells. Overexpression was induced during 24 hours after spheroid formation (Fig. 5a-b). Under the conditions used, overexpression usually occurred in a subset of cells in a spheroid (Fig. S5a). We combined GFP detection using two independent antibodies with detection of primary GFP fluorescence to identify cells that overexpressed a particular protein (overexpressors), the direct neighbors of overexpressors that did not themselves overexpress the protein (neighbors), and non-overexpressing cells that were not neighbors of an overexpressing cell (bystanders) (Fig. 5c). Further, we assigned weakly GFP-positive cells that were localized next to strongly overexpressing cells as ambiguous, since discriminating weak overexpression from spurious positivity due to spatial proximity was not possible. In total, we assessed six replicate spheres for each construct and 30 mock-transfected spheres as technical negative controls. We analyzed more than 500,000 cells from 1,968 spheroid sections from 278 spheroids (Tab. S2).

**Figure 5.**
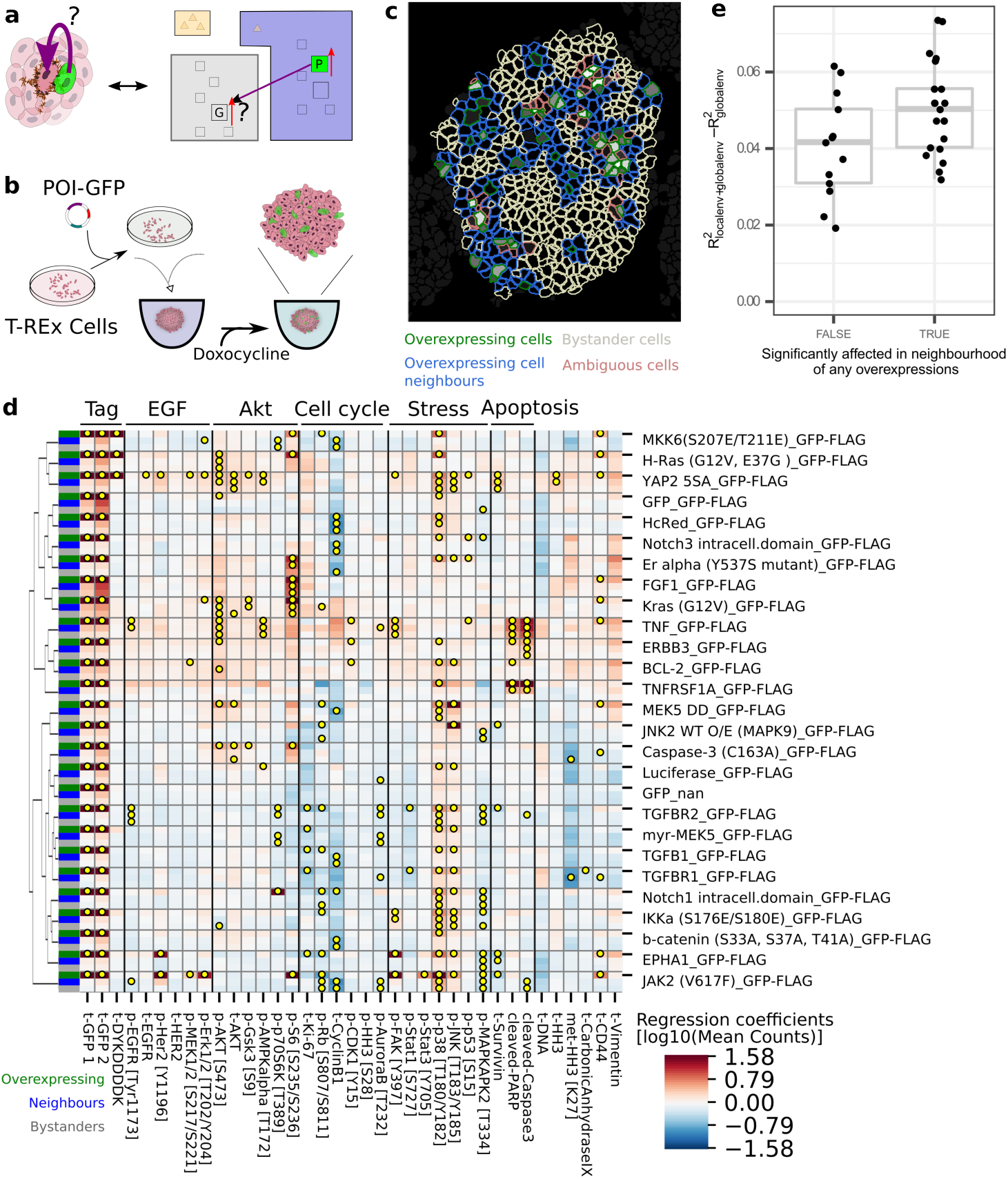
Systematic overexpression reveals spatial effects of signaling deregulation. **a** The schematic (left) depicts a spheroid with a cell overexpressing a construct of interest (green) and a neighboring cell. Protein overexpression could have intracellular as well as neighborhood effects. The illustration (right) depicts an overexpression situation and the question of whether overexpression of protein P (green) alters expression of marker G in a neighboring cell. **b** A schematic of the overexpression system used in this study. Inducible transient transfection leads to GFP-tagged protein overexpression in a fraction of cells in spheroids. **c** A representative spheroid is shown illustrating the identification of overexpressing cells (green), their neighbors (blue), and bystander cells (white). In cells classified as ambiguous (pink) we could not distinguish between overexpression in the cell itself or signal spillover from overexpressing neighbors. **d** Matrix of overexpression estimated effects for constructs (rows) versus effects on markers (columns) classified as intracellular (blue), neighborhood (green), and bystander (gray) in column at the far left. Yellow dots indicate strong, significant effects (p<0.01, q<0.1, fold change > 20%, neighbor/bystander effects: > 0.1x internal effects). All controls and constructs with neighborhood or bystander effects on more than one marker are shown here (see Fig. 5b for data for all constructs). **e** Each dot represents the average dependence of markers on the local neighbourhood after subtracting the dependence on the global environment in unperturbed T-REx-293 spheres grown for 96 hours. Markers significantly affected by in the neighbourhood of any overexpression (TRUE) show on average a higher dependence than markers never affected by any overexpression (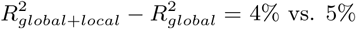 vs. 5%, t-test p = 0.03, Kruskal–Wallis p=0.05, n=34).

For each of the overexpression constructs, we tested whether overexpressor, neighbor, or bystander cells were significantly different in their marker expression from cells of mock-transfected spheres (linear mixed-effect model, p<0.01, q<0.1, fc> 20%, Methods). While intracellular effects would be largely cell-autonomous, we expected that effects on direct neighbors would be dominated by a combination of juxtacrine and paracrine effects. Further, we assumed that bystanders would mainly be affected by longer range paracrine effects of cells in the measured plane as well in the planes above and below the evaluated cell, though juxtacrine effects of off-plane cells could also plausibly contribute to bystander effects.

First, we examined cell-autonomous effects of overexpression. We observed a stress response in overexpressors for most constructs (p-p38: 84%, p-SAPK/JNK: 64%, Fig. 5d, Fig. S5b), including three of the four negative control constructs. A nonspecific stress response to overexpression was not unexpected [40] and thus was not reported as an overexpression-specific effect or included in reported statistics except when explicitly mentioned.

We observed that overexpression of 22 of 32 intracellular signaling proteins, eight of nine ligands, and ten of ten receptors but none of the four negative controls significantly affected more than one intracellular marker (Fig. 5d, Fig. S5b). This indicates that overexpression of most of the constructs perturbed the intracellular state. The overexpression effects were often consistent with known functions of the overexpressed protein and usually involved multiple markers in the relevant pathway (Fig. 5d, Fig. S5b). For example, EGFR overexpression increased total EGFR, p-HER2, and p-ERK1/2 [41](Fig. S5b). FGF receptor overexpression strongly activated its downstream target p-ERK1/2 as well as p-EGFR and p-HER2 [42] (Fig. S5b). TGF beta and TGF beta receptor 3 overexpression both reduced Ki67 and p-RB levels significantly [43](Fig. 5d). Reassuringly, in the three cases, where an antibody in our panel detected the overexpressed protein, we detected the levels of the overexpressed proteins as significantly higher. Where a phosphorylation site in the overexpressed protein was monitored, we observed an increase upon overexpression in three of four cases. The exception was EGFR overexpression, which increased total EGFR and phosphorylation of its interaction partner HER2 but surprisingly did not increase levels of p-EGFR. Overall, these data show that our approach detects biologically expected intracellular responses to overexpression.

We went on to examine the effects of overexpression on neighboring cells using an additional criterion to identify relevant changes: We found that 2.75 *±* 1.73% of the intensity of the signal from an overexpressor, based on GFP, was detected in the neighboring cell (inter quartile range (iqr) [1.6, 3.4], max 8.7), and signal was also detected, although to a lesser degree, 1.0% *±* 1.1%, in bystanders (iqr [0.3, 1.5], max 4.3). Thus, we only considered an increase in marker levels of neighboring or bystander cells significant if the difference was greater than 10% for neighbors or 5% for bystanders of the signal in the overexpressor.

Of the 55 constructs tested, nine caused changes in more than one marker in neighboring cells and ten caused changes in bystander cells (Fig. 5d). Notably, all of these are part of the set of 40 constructs that caused specific and significant changes in internal cell state. In contrast, apart from myristoylated MEK5, none of the constructs without cell autonomous effects exhibited effects on more than one marker in neighbours or bystanders. Of the constructs that caused changes in expression of at least two markers in neighboring cells, one was a ligand (of eight tested), five were intracellular signaling proteins, and four were receptors (of 10 tested) (Fig. 5d). The effects on neighbors (marker fold change iqr 1.2-1.5, max 2.4) and bystanders (iqr 1.2-1.4, max 2.0) were usually weaker than internal effects of overexpression (iqr 1.3-4.9, max 42.8).

Of the intracellularly affected markers, 18% were also significantly changed in neighbors and 13% were altered in bystanders. This is more than ten times more frequent than markers not intracellularly changed. In cases, where a marker was increased both in overexpressors and in neighbors, the effect on neighbors was an average of 77 *±* 29% of the intracellular effect (min 22%, max 132%). Similar values were observed for bystander effects.

We next asked whether markers that depended on the neighborhood in unperturbed 293 spheroids were more likely to be perturbed by overexpression in neighboring cells. We found that the set of markers (57% of all markers) that were significantly perturbed in either neighbors or bystanders in any of the overexpression experiments indeed had a slightly higher dependence on neighboring cell marker levels in our analysis of unperturbed T-REx-293 spheres grown for 96 hours (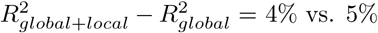, t-test p = 0.03, Kruskal–Wallis p=0.05, n=34, Fig. 5e).

For example, overexpression of the ligand TNF and its receptor TNFRSF1A caused significant increases in apoptosis markers in overexpressors, their neighbors, and, in the case of TNF, in bystander cells (Fig. S5c). This neighbor effect is consistent with our observations in unperturbed spheroids, where we found that the apoptosis marker cleaved PARP was dependent on the neighborhood in large T-REx-293 spheroids (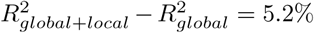 Fig. 4c). We also found that markers of AKT signaling (pAKT, pGSK, and pS6), the three markers most strongly linked to the cellular neighborhood in the variability analysis 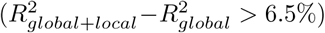), are also affected in neighbors of cells overexpressing constructs such as hyperactivated YAP 5SA or hyperactive KRAS G12V, suggesting that AKT signaling may not only be regulated by environmental factors but also via direct cellular interactions. In summary, these perturbation experiments allowed us to detect non-cell autonomous effects of overexpression on neighboring cells. Markers that showed a dependency on cellular neighborhood in steady-state experiments tended to be more likely to be modulated by these perturbations in neighboring cells.

## 4 Discussion

### 4.1 Quantification of environmental influence on cellular phenotypes

We coupled a 3D spheroid tissue model system with highly multiplexed imaging to characterize the influence of global and local cellular environment on cellular phenotypes.This allowed us to observe that measures of local and global environments and internal cell state are not independent in their abilities to predict marker variation. Rather, there were strong non-additive interdependencies among these factors. Specifically, and consistent with the spatial architecture of spheroids, measurements of the local neighborhood of a cell captured marker variability explained by the global environment. Moreover, spatial autocorrelation alone explained much of the marker variation captured by local and global environmental effects, and comprehensively measured intracellular state markers (including cell-cycle markers) recapitulated much of the explanatory power of all environmental effects combined.

Such interdependencies must be taken into account in studies aiming to deconvolve the contributions of environmental factors to phenotypic variability. Although a comprehensive intracellular marker measurement allows to predict the behavior of an environmentally sensitive marker even without an environmental measure, this does not mean that such environmental effects do not exist. In fact, they could even be the causal reason for the behavior of the marker. Such effects could be missed if the interdependence between explanatory factors is not taken into account. Intracellular markers have been validated as accurate surrogates for environmental conditions in non-spatial cytometry analyses [44]. Examples are hypoxia markers as surrogates for cell position in an oxygen gradient and phosphorylated receptor levels as surrogates for ligand binding.

Interdependencies in spatial measurements have been acknowledged in *E. coli* [45]. However, a recent approach developed for multiplexed data analysis, Spatial Variance Component Analysis, assumes that contributions of spatial proximity (environment), neighborhood levels (cell-cell interactions), and cell state (intrinsic) are independently additive [46]. This assumption may bias results. To account for interdependencies in our own dataset, we used a step-wise regression approach, in which predictors were added in increasing order of explanatory power. Additionally, we split internal markers into those that are cell-cycle-related and those not involved in the cell-cycle to account for the known importance of cell-cycle in marker variation [32–34, 47]. We showed that this regression approach was able to quantify how phenotypic markers depend on the influence of the global environment, local neighborhood, autocorrelation and internal markers. We confirmed several of the identified patterns by visual inspection of images. Our simple model system allowed us to compare the interdependent effects of environmental and cell autonomous factors on cellular phenotype in different cell lines and growth conditions and to quantify marker-specific differences in spheroid organization.

Around 50% up to 85% of marker variability within homogenous 3D spheroid sections could be explained using our complete linear model, which included global environment, local neighborhood, local autocorrelation and internal cell state modules. For low-abundance markers (< 1 average count per cell pixel), technical detection noise likely dominated marker variability. This was not the case for markers expressed at higher levels, however. An additional technical source of variability could be that we are relying on 6 *µm* thick slices of cells, measured at a lateral resolution of 1×1 *µm*. At this resolution, pixels may belong to more than one cell, and segmentation is unlikely to be perfect, which introduces technical variability. Further, our readouts do not represent full cells but random, 6 *µm* thick slices through cells, which could introduce technical sampling variability, in particular when markers are not uniformly distributed across the cell. Finally, our analysis assumes linear marker relationships, which may explain the lack of fit of our model to some extend.

### 4.2 Challenges in adapting the analysis to complex tissues

A future key challenge will be to extend our and similar approaches to heterocellular tissues, which are more representative models of biological systems than the spheroids analyzed here. Such tissues are likely to be highly structured with different cell types confined to specific locations, resulting in strong cell-type co-occurrence patterns. Applying methods that quantify relationships between cells and their neighborhood agnostic of cell types will likely capture cell-type co-occurrences as neighborhood effects [46]. Although meaningful, co-occurrence of cell types does not provide the full picture of how the cellular phenotype is influenced by the environment or neighborhood. Lineaging approaches could help mitigate the confounding effects of co-occurring cell types and provide a ground truth for phenotypically comparable cells.

Furthermore, identification of biologically relevant spatial gradients in complex models will be much more challenging than in our symmetrical spheroid model, which allowed estimating these gradients based on the known location of the source (i.e., nutrients in the medium) and of the sink (i.e., cells). Gradient characterization will be important, as illustrated here we showed for spheroids where a readout for such gradients is key to understanding the cause of observed spatial variability and correlations. While it may be theoretically possible to estimate the number of relevant biological gradients in complex tissue based only on phenotypic information [48], capturing quantitative information on these gradients will require a stereotypical tissue structure and biological domain knowledge. Identifying gradients in tissues and using them as biologically relevant coordinate systems will aid in identification of causes of phenotypic variability and will enable comparisons across tissue samples.

### 4.3 The influence of extreme cell states on neighboring cells

We used a chimeric overexpression system to systematically assess the effects of deregulated signaling on cellular neighborhoods in the spheroids formed by T-REx-293 cells. Of the 55 constructs overexpressed, 73% induced intracellular changes in multiple markers, and around 20% caused non-cell autonomous effects on neighboring and bystander cells. This indicates that chronic overexpression of signaling proteins alters not only the intracellular state of the cell overexpressing the signaling protein but also, at least in some instances, cell states of neighbors.

It is likely that our analysis missed some effects: Although our marker panel covers multiple signaling pathways and cellular processes, our previous studies have shown that overexpression can alter signaling transiently, without an effect on steady-state marker levels at the time of measurement [49, 50]. Focusing on steady-state levels in our analysis meant that we missed such dynamic effects. Further, misfolding and mislocalization of tagged, overexpressed constructs can lead to non-physiological effects, including the stereotypic intracellular stress responses evident in our data. However, we also observed clear construct-specific intracellular responses that were consistent with known biological functions of the overexpressed protein.

Our use of linear mixed effect models assumes normality, heteroscedasticity, and spatial independence of residuals. These assumptions are violated to various degrees, potentially leading to false positives and false negatives. Moreover, our analysis required dichotomizing the data into overexpressing and non-transfected cells. Even in non-transfected cells, background signals are generated by anti-GFP antibodies, likely due to non-specific binding. Further, fluctuations in signal detection sensitivity through weak staining could lead to false negatives. We used a variety of approaches to maximize the reliability of our results. We sought to increase sensitivity and specificity by aligning our two antibody-based IMC GFP readouts with fluorescent microscopy images of primary GFP fluorescence. We also trained a pixel classifier to identify ‘positive’ pixels leveraging the information from all available channels. In addition, we introduced a quality control step to identify tissue folds, a common source of elevated background staining, to decrease false positives. We further used stringent cutoffs on p-values, false discovery rates, and fold changes and used our negative control constructs (two different GFP constructs, HcRed and luciferase) as additional guidance. None of these controls caused significant changes in more than one marker apart from the stereotypic stress responses observed in overexpressors, neighbors, and bystanders. In contrast, 80% of overexpressed ligands, receptors, and cancer-related signaling proteins showed multiple responses in overexpressors and sometimes in neighbors. From this we conclude that our approach reliably captures how cells grown as spheroids respond to intracellular overexpression as well to overexpression in cells in their environment.

## 5 Conclusion

In conclusion, we developed a novel tissue barcoding workflow for simultaneous processing of up to 240 microtissues and used this setup to generate a large multiplexed imaging dataset of homogeneous 3D spheroids with single-cell resolution. Our dataset will be a useful resource for the further development of algorithmic approaches describing spatial variability in cellular phenotypes. We have assessed how cell state as well as local and global environment affect cellular phenotype and report hierarchical interdependencies of these factors in their ability to explain marker expression. Our approach is broadly applicable and will enable the robust characterization of more complex tissues, from co-cultures to heterocellular organoids and small embryos. We also demonstrated that our approach is compatible with perturbation studies and identified cell autonomous and neighborhood effects of overexpressed cancer-related signaling proteins. We envision that this approach could be used to systematically study the impact of perturbations on the organization of simple and complex microtissues. This strategy could, for example, provide insight into how drug treatment alter the interplay of cell types in healthy and diseased tissue.

## 6 Materials and Methods

### 6.1 General

#### 6.1.1 Cell lines

T-REx-293 cells (Invitrogen) and DLD1 cells (Flp-In T-Rex DLD-1, donation Stephen Taylor lab, University of Manchester) were grown in high-glucose DMEM (D5671, Sigma). HT29 (ATCC HTB-38) and T47D (ATCC HTB-133) cells were grown in RPMI-1640 medium (R0883, Sigma). The media were supplemented with 100 *U/ml* penicillin, 100 *mg/ml* streptomycin and 2 *mM* L-glutamine (GIBCO, Invitrogen). For T47D cells, 0.2 *U/ml* human insulin were added. All media were filtered through a 0.2 *µm* membrane (Nalgene, Thermo). 1x TrypLE Express (Life Technologies) was used for cell passaging and harvesting. Mycoplasma was not detected with a MycoAlert PLUS Mycoplasma Detection Kit (Lonza) in any of the cell lines.

#### 6.1.2 Antibody conjugation

Isotope-labeled antibodies were prepared using the manufacturer’s standard protocol using the MaxPAR antibody conjugation kit (Fluidigm). Conjugated antibody yield was determined based on absorbance at 280 *nm*. For long-term storage, antibodies were stored at 4^°^*C* in PBS Antibody Stabilization solution (Candor).

#### 6.1.3 Spheroid cultivation

Spheres were grown in ultra-low adhesion 96-well Spheroid Microplates (Corning) in a volume of 100 *µl*, covered by an breathable membrane (Breathe Easier, Diversified Biotech, BERM-2000) and incubated in a 5% CO_2_ incubator at 37^°^*C*. Spheres were fixed by adding 30 *µl* of 16% PFA per well, incubated with shaking at 200 *RPM* for 5 *min* and then stored overnight at 4^°^*C*. The next morning, the spheres were washed four times with 150 *ul* 1x PBS (pH 7.4) using a Biomek Fx robot (Beckmann Coulter).

#### 6.1.4 Bright field imaging

Bright field imaging of intact spheres was performed using an ImageXpress Micro XL Widefield High Content Imaging microscope (Molecular Devices, 4x objective, NA 0.20) at multiple z-planes. Spheres were imaged 2 *h* before PFA fixation as well as after PBS washing the next morning. Plates were acquired twice, rotating the plate by 180^°^ to avoid imaging artifacts.

#### 6.1.5 Barcoded spheroid assay

Barcoding reagents were added using a redundant barcoding scheme (60-well: [17], 120-well: [50]) and plates were incubated for 1 *h* shaking at 200 *RPM*. Spheres were washed four times with 150 *µl* 1*x* cell staining medium (CSM, PBS (pH 7.4) 0.5% BSA). Spheres were pooled into 6 *ml* collection tubes (pre-incubated with CSM) using 200 *µl*-wide bore tips (FX-255-WB-R, Corning Axygenx). All pipetting steps were implemented with a Biomek Fx robot (Beckmann Coulter).

The pooled spheres were washed twice with 4*ml* PBS; samples were centrifuged 1 *min* at 100 *xg* after each wash. Then, 1 *µM* monoisotopic cisplatin was added in 1 *ml* PBS, and the tubes were incubated for 30 *min* with shaking at 200 *RPM*. The spheres were washed twice with 4 *ml* CSM, and all but 200 *µl* of supernatant was removed. Spheres were incubated for 5 *min* at 37^°^*C*, then 4 *ml* 12% gelatin (white-gold extra, Dr. Oetker, swell cold for 10 *min*, then dissolve for >4 *h* at 60^°^*C*, keep at 37^°^*C*) in 0.1 *M* PBS was added). Spheres were incubated for at least 10 *min* until they sunk to the bottom. A cylindrical mold with a bottom diameter of 2.5 *mm* was prepared by positioning a glass rod in 12% agarose in a 1.5 *ml* tube. After solidification, the agarose mold was incubated with gelatin at 37^°^*C*. The spheroids were transferred into the agarose mold in multiple steps. Using a centrifuge at 200 *xg* at 37^°^*C*, spheres were spun down in the mold until there was an even layer at the bottom of the cylinder. Manipulation using a pre-warmed pipette tip was used to improve the positioning. Once spheroid positioning was even, the mold was cooled at 4^°^*C* overnight. Then, the agarose mold was carefully broken open and the spheroid gelatin plug was incubated for 1 *h* first in 15% sucrose (Sigma), then for 4 *h* in 30% sucrose in doubly distilled H _2_O with 0.004% trypan blue (Sigma). The pellet was positioned into OCT medium (Tissue-Tek) and frozen in -40^°^*C* isopentane (Sigma) and stored at -80^°^*C*. The OCT block was cut using a cryo-microtome (thickness: 6 *µm*, object/knife temperature -17/-15^°^*C*), sections were transferred to microscopy slides (Superfrost Plus, Thermo Scientific), dried overnight at room temperature, photographed, and stored at -80^°^*C*.

#### 6.1.6 Antibody staining

Sections with minimal tearing covering the whole volume of the pellet were selected. Cuts were transferred from -80^°^*C* into TBS (50 *mM* Trizma base (Sigma), 50 *mM* NaCl (Sigma), *pH* 7.6) and washed three times with TBS (10 *min* each wash). Individual sections were marked with a Dako Pen (Agilent) and blocked with 3% bovine serum albumin in TBS-T (TBS+ 0.1% Tween 20 (Sigma)) for 1 *h*. The antibody solutions were added (Table S1). A spillover slide was created for the whole panel by spotting 0.3 *µl* antibody in 0.5 *µl* 0.4% tryphan blue on an agarose-coated slide [51]. Then, the blocking buffer was removed and each section stained with 12 *µl* antibody mix overnight at 4^°^*C* in a hybridisation chamber. Next day, the slides were washed 3x in TBS for 10 *min*. 20 *µl* per site of 1 *µM* Iridium Intercalator (Fluidigm) was added for 10 *min*. Then the sites were washed with TBS and 1 *µM* Hoechst 33342was added for 6 *min*. Finally, slides were washed 3x 10 *min* in TBS, dipped 3 *s* into ddH _2_O and blow dried with compressed air.

#### 6.1.7 Fluorescent imaging

The dried slides were imaged using a Axioscan Slide Scanner Z1 (Zeiss) using the DAPI as well as the GFP channel, where appropriate.

#### 6.1.8 Imaging mass cytometry

Slides were ablated using a Helios Imaging Mass Cytometer (Fluidigm) at nominal resolution of 1 *µm*^2^ and an ablation frequency of 400 *Hz*.

### 6.2 Cell line physiology

For the unperturbed experiments, cells were seeded into the spheroid microplates at concentrations of 1x, 0.5x, and 0.25x, where the 1x concentrations were 3,200 cells per well for T-REx-293 cells, 6,400 cells per well for DLD1 cells, 2,000 cells per well for T47D cells, and 2,000 cells per well for HT29 cells. Cells were grown in five replicates and each plate was barcoded using a 60-well barcoding scheme [17]. One plate, p173, was fixed and barcoded after 72 *h*, and the other, p176, was fixed and barcoded after 96 *h*. Monoisotopic cisplatin (194Pt, 1 *µM*) was added after pooling the spheroids. For the 72 *h* timepoint, data was acquired on 18 cryo-sections, and for the 96 *h* timepoint data was acquired on 16 cryo-sections.

### 6.3 Chimeric overexpression experiments

#### 6.3.1 Constructs

We used a library generated from the entry clones of a previously published cancer signaling constructs library [39]. We added constructs encoding biologically relevant ligands and receptors from the human ORFeome V8.1 library (Dharmacon) via NEXUS Personalized Health Technologies at ETH Zurich [52]. Destination vectors, including pDEST, pcDNA5 FRT TO-eGFP, and pDEST 3’ Triple Flag pcDNA5 FRT TO, were kindly provided by Anne-Claude Gingras (Lunenfeld-Tanenbaum Research Institute, Toronto, Canada [53]). Tagged expression vectors were generated via Gateway Cloning. Sanger sequencing was used to confirm the clone identity before transfection. Constructs were arranged on a master plate in a randomized fashion with control wells evenly distributed over the plate.

#### 6.3.2 Experiment

T-REX 293 cells were seeded at a density of 20,000 cells per well in 100 *µl* medium into two 96-well flat-bottom cell culture plates (p155 and p156), using the normal medium prepared with tetracycline-free FBS (S182T-500, Biowest). After 24 *h* incubation, transfection was done using the JetPrime transfection system (Polypus) according to the manufacturer’s instructions: For each construct a master-transfection mix of 22.5 *µl* jetPrime buffer, 0.5 *µl* jetPrime reagent, and 2.5 *µl* of 0.1 *µg/µl* DNA was prepared. An aliquot of 10 *µl* of this master mix was added dropwise to each well. After 5 *h*, the cell culture medium was changed using the Biomek robot under semi-sterile conditions.

After 24 *h*, the cells were washed with PBS and detached with 100 *µl* 10x TrypLE Select Enzyme (Gibco) per well. Cells were resuspended in 100 *µl* medium. From these plates, cells were distributed into the spheroid microplates: From plate p155, 4 *µl* cell suspension per well was added to each well of plates p161 and p163. From plate p156, 20 *µl* of mock-transfected cells from border wells were transferred to each well, before 2 *µl* of the suspension was added per well to plates p165 and p167 and 4 *µl* was added per well to plates p169 and p171.

After 48 *h*, the spheres were imaged with bright field microscopy and 2 *µl* of 50 *µg/ml* tetracycline hydrochloride (Sigma) in PBS to a final concentration of 1 *µg/ml* was added to each well. After 24 *h*, spheres were fixed and barcoded. For barcoding, three pairs of plates were barcoded using the 120-well barcoding plate layout [50] (Plate 1: p161, p165, p171, Plate 2: p163, p167, p169). Then plates p165 and p171 and plates p161 and 163 were pooled, and cisplatin (194Pt) was added. Plates p167 and p169 were pooled, and cisplatin 198Pt was added. Finally, plates p165, p171, p167, and p169 were pooled into one spheroid plug with 240 wells. p161 and p163 were embedded as a spheroid plug of 120 wells. After sectioning, 20 slices of the 120 well plug and 48 slices of the 240 well plug were selected for staining.

## 7 Analysis

A repository containing all raw data and a reproducible workflow to generate all analyses from raw data is in preparation.

### 7.1 Bright field Image Analysis

#### 7.1.1 Quantification

In order to robustly determine spheroid diameter, we used a pipeline based on supervised pixel classification by Ilastik [54]; this process identified spheres despite intensity variations. We used CellProfiler for segmentation and quantification [54].

#### 7.1.2 Quality control

As quality control, bright field images of each well were manually screened for spheroids with growth defects, such as particle or fiber inclusions, blinded for the spheroid growth condition.

### 7.2 General spheroid IMC image analysis

#### 7.2.1 Quantification

To robustly identify spheroids in IMC images, we used supervised pixel classification by Ilastik [54] to identify spheroid centers, borders, and background, and used CellProfiler [54] to segment the resulting probability maps. To identify cells, we used similar approach, using Ilastik to classify pixels into nuclear, cytoplasmic/membrane and background and CellProfiler to identify cells based on the resulting probability maps. Within cell regions we quantified marker levels, applied compensation [55], and calculated other spatial features. Neighbors were identified by expanding each cell object by 3 pixels and identifying touching cells. We built an analysis framework based on Python [56], SQLite, and Scanpy [57] to handle and analyze the data.

#### 7.2.2 IMC - fluorescent slide scan imaging alignment

Fluorescent slidescan images and IMC acquisitions were aligned with a fully automated iterative alignment process using Trakem2 [58]. For the slidescan images, the DAPI channel and for the IMC acquisitions, the Iridium channel was used. First, whole spheroid plug sections were coarsely aligned using a rigid transform estimated from the machine provided global coordinate system. Then, the sections were aligned using a rigid alignment estimated by TrakEM2. Finally, individual cropped spheroid sections images of the two modalities were fine-aligned using rigid alignment by TrakEM2.

#### 7.2.3 Quality control

On a cellular segmentation level, several quality control criteria were applied: Sphere membership: At least 95% of all pixels of a cell need to be within the sphere segmentation region. Sphere ambiguous cells: Cells closer than 20 pixels to any other sphere in the image were excluded as they might be ambiguous.

- Cell size: Cells smaller than 10 pixels were excluded.
- Border cells: Cells directly touching the outer spheroid segmentation border were excluded.
- Main sphere: Cells not belonging to the largest contiguous cell mass of still valid cells were excluded.
- Fold classifier: A pixel classifier was trained in Ilastik based on DAPI fluorescence to identify folded areas in the spheres.

On a sphere slice (image) level, we used the following criteria to exclude images with all their cells:

- Not small: sphere slices with less than 10 valid cells (above) were excluded.
- Manual QC: Quality control image consisting of raw images, quantification, spheroid and cell segmentation of DNA (193Ir), Histone H3, and 194Pt, and 198Pt channels were visualized for each spheroid slice with an anonymized ID. This allowed the visual identification of image artifacts such as folds, bubbles, tissue tearing, and spheroid mis-segmentations blind to spheroid growth conditions.

Additionally, we used bright field images to identify wells with particles, fibers, deformed spheres as well as multiple spheres. Sphere slices from these wells were excluded.

The effect of the individual QC steps on the two datasets can be found in table S2.

#### 7.2.4 Data transformation

If not otherwise indicated, the spillover-compensated mean pixel intensity per cell area was used as a readout. The data was *log*10(*x* + 0.1) transformed and winsorized using the 0.1th percentile.

#### 7.2.5 Debarcoding

For debarcoding, cells within 30 pixels from the outer spheroid border were considered. Over all images belonging to a spheroid plug, each barcoding channel was binarized with the average barcode channel intensity. Then, for each spheroid, the number of valid barcodes was determined. Due to the robustness of the barcode schemes used, false positives were not expected to be frequent. Sphere slices were assigned to the most common valid barcode in the slice. As quality control, at least 10 cells were required to be assigned to this barcode, and the most common barcode was required to be at least twice as frequent then the second highest barcode. Images without valid identification were excluded from further analysis.

#### 7.2.6 Distance to border correction

Distance to the border of the spheroid slices can overestimate distance to border in the spheroid, as they represent spherical segments at different heights of the sphere. As the real spheroid diameter can be estimated by bright field imaging, and assuming that spheroids are indeed spherical, this was correcting by the formula:

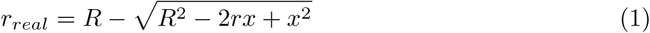

where *r*_*real*_ is the real distance to the sphere border, R is the radius of the sphere measured in the bright field images, *r* is the radius of the segment, and *x* is the measured (non-corrected) distance to the border in the segment.

#### 7.2.7 UMAP and cluster analysis

Uniform Manifold Approximation and Projection (UMAP) [59] and clustering via the Leiden algorithm [60] were performed via SCANPY [57].

### 7.3 Marker variability analysis

For the marker variability analysis, the level of each marker was predicted by a linear model (Fig. 3a):

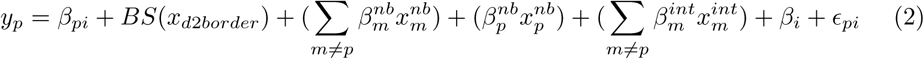

where:

- *y*_*p*_: the level of marker *p* in a cell
- *β*_*pi*_: technical staining/batch effect for marker *p* of the image *i* that the cell is part of
- *BS*(*x*_*d*2*border*_): a nonlinear function of distance to border (*x*_*d*2*border*_ represented by a polynomial B-spline of degree 3 with 10 knots distributed located at the deciles (10 quantiles)
- 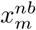: average levels of marker *m* in direct neighbouring cells
- 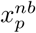: average levels of predicted marker *p* in direct neighbouring cells
- 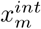: cell internal marker levels of marker *m*

The models and submodels were fitted using the statsmodels library [61].

If not mentioned otherwise, the reported variability explained (*R*2) for each model was the adjusted *R*2 relative to the adjusted *R*2_*tech*_ - the *R*2 of a model only containing an image-specific intercept. This prevented variability in signal differences resulting from technical issues (e.g., due to staining or acquisition) from being attributed to biological variability. This corrected *R*2 is calculated according to:

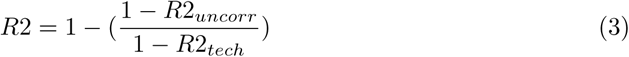

For the correlation heatmaps and distance to border plots (Fig. 2c, S3), an image-specific intercept was fit. This intercept was subtracted before calculating the Pearson correlation.

#### 7.3.1 Permutation analysis

We fit all possible sequences of adding the modules for global environment (*BS*(*x*_*d*2*border*_)), local environment 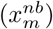, autocorrelation 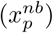 and cell state 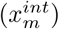 to the model, for each marker and condition, and recorded the additional marker variability explained at each step in each sequence. For each sequence, we calculated the variance of the additional variability explained by each added module. Given that the total variability explained is independent of the sequence, high variance suggests that marker variability is explained by a few modules and low variance suggests that marker variability is explained by multiple modules. In a strictly hierarchical dependency structure, modules higher in the hierarchy should contain the variability explained by those lower in the hierarchy. Adding a more explanatory module before a less explanatory one, will lead to the former fully capturing the variability of the latter, yielding high variance. Sequentially fitting modules in line with the hierarchy of explanatory power should maximise the contributions of each module, thus reducing the variance. The optimal order would correspond to the sequence with least variance.

### 7.4 Chimeric overexpression analysis

We trained a pixel classifier based on the two IMC GFP antibodies as well as the aligned primary GFP fluorescent slidescan image to robustly detect overexpressing image regions. We calculated the average pixel-wise probability for overexpression and required this to be more than 0.01 (estimated false discovery rate: 0.003) for a cell to be classified as ‘overexpressing’. Cells with an average pixel-wise probability higher than 0.01 but lower than 30% of the maximal value observed in neighbouring cells were added to an ‘ambiguous’ category. Other cells within 6 pixels of ‘overexpressing’ cells were classified as ‘neighbour’ cells. All other cells were classified as ‘bystanders’. This classification was not reliably possible for spheres transfected with a FLAG-only construct without GFP, due to FLAG antibody background staining. All cells in such spheres were classified as ‘bystander’ cells.

We used a linear mixed effects model to estimate marker levels independently of the effect of belonging to the overexpressing, neighboring, or bystander cell class of a specific construct. The following model was used:

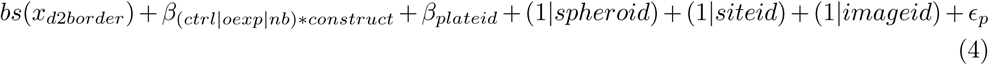

Where :

- *y*_*p*_: the predicted marker level
- *BS*(*x*_*d*2*border*_): a nonlinear function of distance to border (*x*_*d*2*border*_: represented by a polynomial B-spline of degree 3 with 10 knots distributed located at the deciles (10 quantiles)
- *β*_(*ctrl*|*oexp*|*nb*)+*construct*_: an intercept for combination of construct and overexpression class
- *β*_*plateid*_: a fixed effect intercept for belonging to any of the 6 plates. This accounts for plate wise effects.
- (1|*spheroid*): a random effect acknowledging that cells from the same sphere are correlated
- (1|*siteid*): a random effect acknowledging that sphere sections/images that were stained and acquired together are not independent, e.g. through staining effects
- (1|*imageid*): a random effect acknowledging that cells from same sphere slide/image are not independent.
- *ϵ* : residual variation. This is assumed to be homoscedasticity.

One model over all constructs was fitted using the lme4 package using the maximum likelyhood estimation [62]. The lmertest package was used to calculate p-values using the Satterthwaite approximation [63, 64]. Multiple testing correction was performed using the Benjamini/Hochberg approach [65].

## 8 Acknowledgements

The authors acknowledge the assistance and support of the Center for Microscopy and Image Analysis, University of Zurich, in particular José Mar í a Mateos Melero, as well as Claudia Meyer from the Institute of Anatomy, University of Zurich, for their expertise and equipment for histological embedding and cutting. We are also in depth for Min Lu and Kris C. Wood for sharing the entry clones for the cancer related signaling constructs library. We are also grateful for all contributors to the open source software packages that made this project feasible. Finally we would like to thank all members of the Bodenmiller and Greber lab for their support and input. BB’s research was supported by a SNSF Assistant Professorship grant, a NIH grant (UC4 DK108132), and by the European Research Council (ERC) under the European Union’s Seventh Framework Program (FP/2007-2013)/ERC Grant Agreement n. 336921.

## 9 Author contributions

Vito Zanotelli (VZ) and Bernd Bodenmiller (BB) conceptualized this study. VZ and Matthias Leutenegger (ML) developed the wet lab and computational workflow with support from BB, Xiaokang Lun (XL) and Fanny Georgi (FG). VZ analysed the data and generated the figures with feedback from BB and Natalie de Souza (NS). VZ, NS and BB wrote the manuscript and XL, FG and ML provided feedback. BB acquired the funding.

## 12 Figures

**Table S1.**
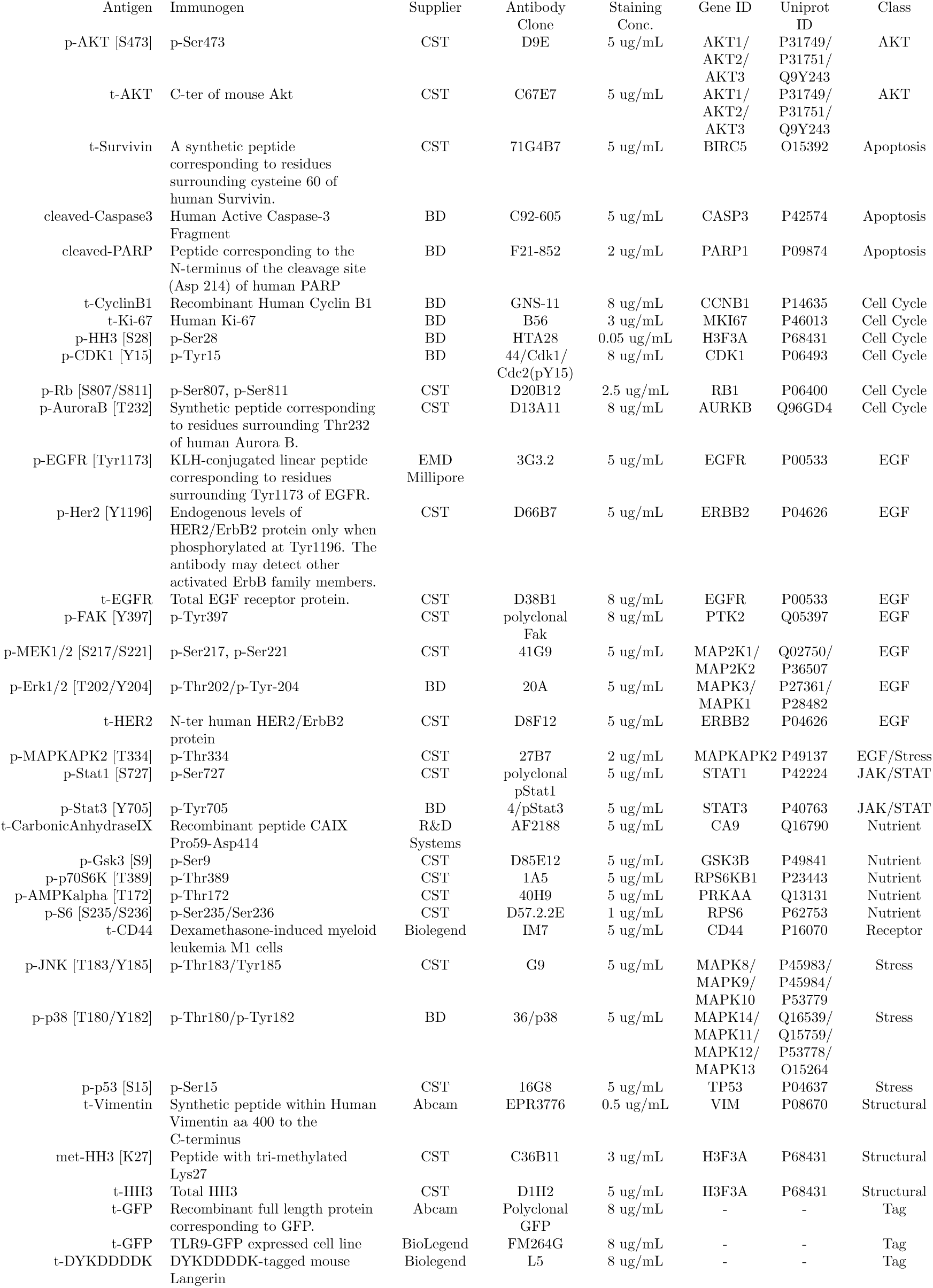
Antibody panel used.

**Figure S1.**
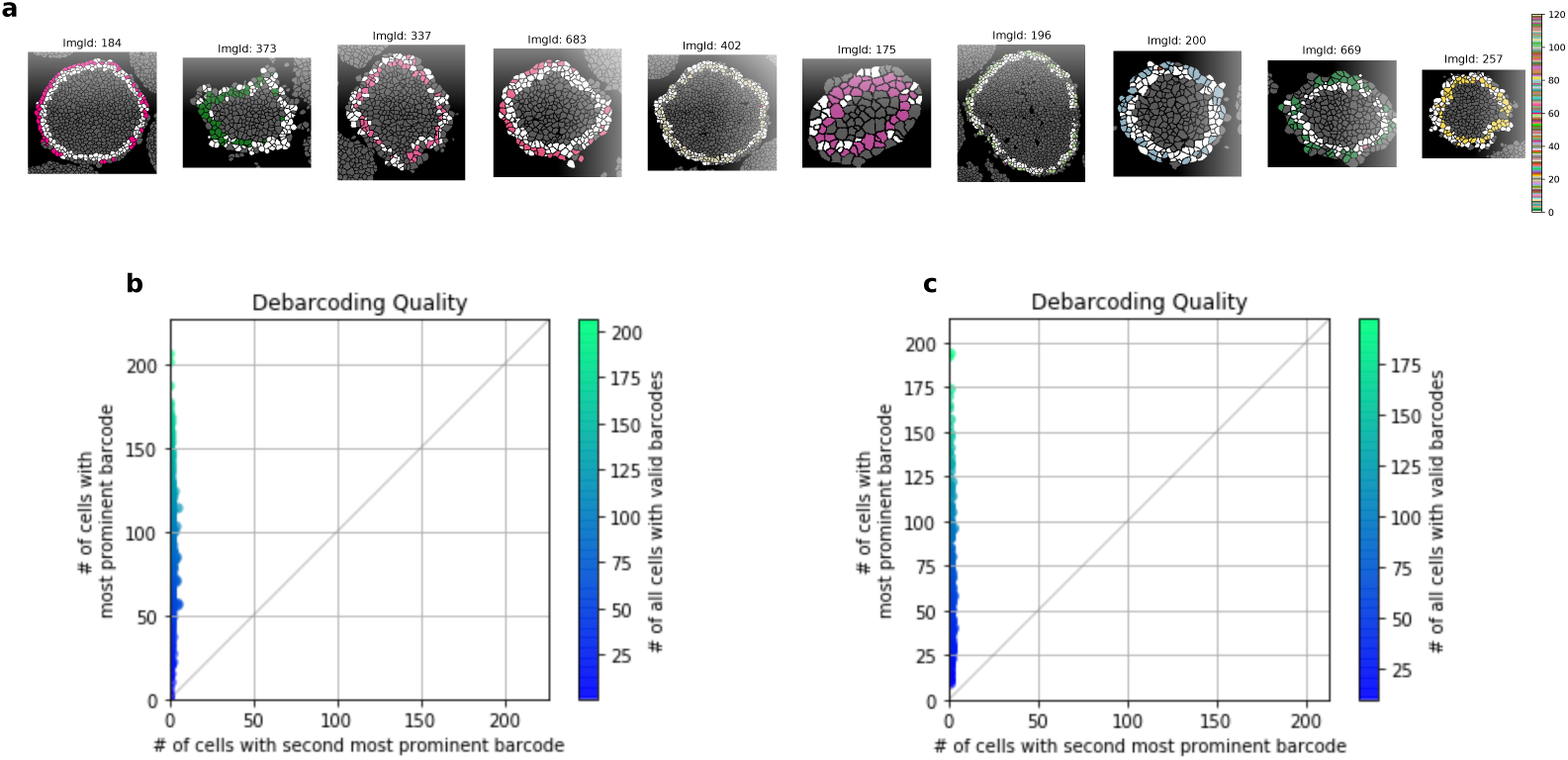
**a** Representative barcoding results on the cell level. Cells were debarcoded by determining if the threshold barcode channels corresponded to a valid barcode (colored cells) or not (white cells). Cells outside of the border of spheres were not considered for debarcoding (grey cells). The sphere section barcode assignment required that the majority barcode was present in at least ten cells and in more than twice the number of cells than the second most abundant barcode. **b** Number of cells with the most prevalent barcode plotted versus the number of cells with the second-most prevalent barcode per sphere from a dataset of 120 barcoded shperes. **c** Number of cells with the most prevalent barcode plotted versus the number of cells with the second-most prevalent barcode per sphere from a dataset of 360 barcoded spheres.

**Table S2.** Table indicating the number of cell slices/ sphere slices and spheres lost at each quality control step for the 4 cell line steady state dataset as well as the overexpression experiment.

| Level | Name | Steady state |  | Overexpression |  |
| --- | --- | --- | --- | --- | --- |
|  |  | Remaining | Removed | Remaining | Removed |
| Cell Slices | Initial number of objects | 349968 |  | 1016872 |  |
|  | Sphere membership | 302626 | 47342 | 848191 | 168681 |
|  | Sphere ambiguous cells | 297053 | 5573 | 826413 | 21778 |
|  | Cell size | 295558 | 1495 | 821340 | 5073 |
|  | Border Cells | 287265 | 8293 | 774453 | 46887 |
|  | Main sphere | 287038 | 227 | 772124 | 2329 |
|  | Fold classifier | 264775 | 22263 | 689671 | 82453 |
|  | Sphere Slice/Spheres QC | 228740 | 36035 | 591009 | 98662 |
| Sphere Slices | Initial number of sphere slices | 736 |  | 2683 |  |
|  | Not small | 668 | 68 | 2515 | 168 |
|  | Manual QC | 576 | 92 | 2096 | 419 |
|  | Debarcoding | 574 | 2 | 2089 | 7 |
|  | Brightfield QC | 517 | 57 | 1968 | 121 |
| Spheres | Initial number of spheres | 120 |  | 360 |  |
|  | Debarcoding | 116 | 4 | 300 | 60 |
|  | Brightfield QC | 100 | 16 | 278 | 22 |

**Figure S2.**
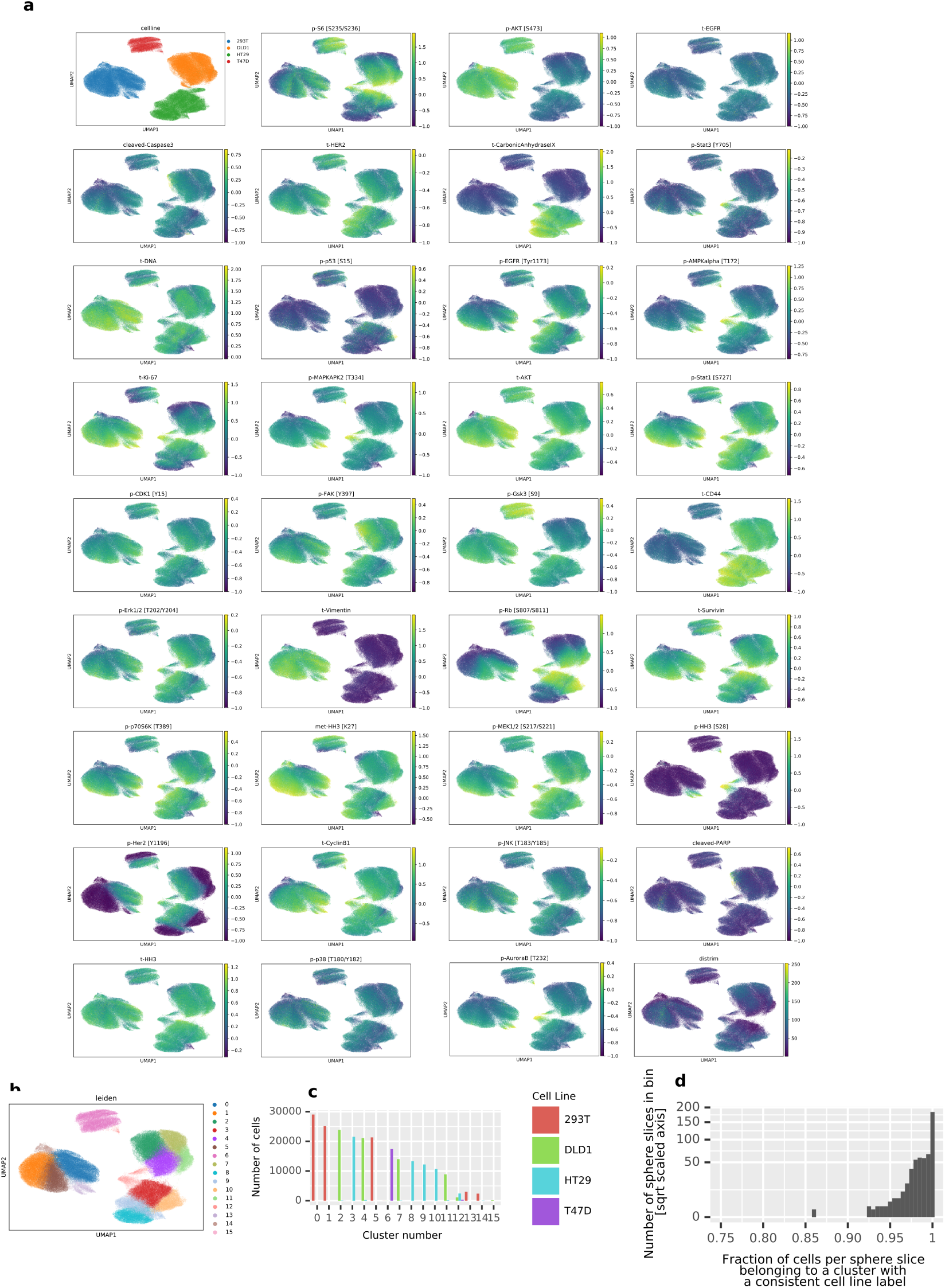
**a** Nonlinear dimensionality reduction with UMAP [59] visualizes average marker levels across the unperturbed cell line dataset. The first graph shows cell line labels assigned by debarcoding. The other panels show marker levels with colors indicative of *log*10(*MeanIntensity* + 0.1) per marker. **b** Clustering of the cell slice data using Leiden clustering [60]. **c** Number of cells from each cell line per cluster. **d** Fraction of cells per sphere slice where the debarcoded sphere slice label was identical to the most abundant cell line label present in the cluster.

**Figure S3.**
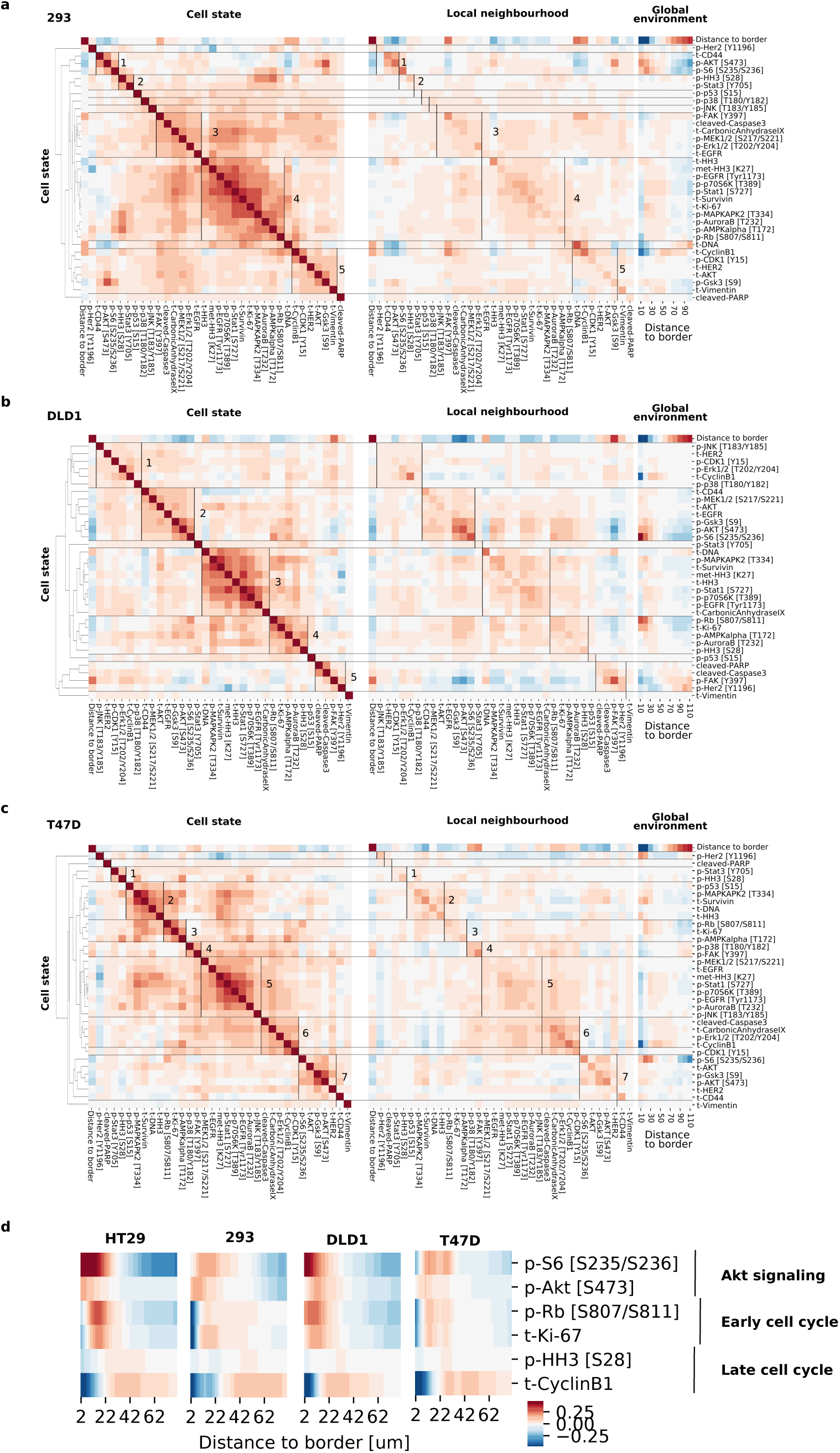
**a-c** Correlation heatmaps as in figure 2d for all cell lines. **d**: Selected growth signaling and cell cycle markers plotted as a function of distance to border for spheres of all cell lines (96 h growth, ca 200*µm* diameter). Scale: centered *log*10(*x* + 0.1) intensity.

**Figure S4.**
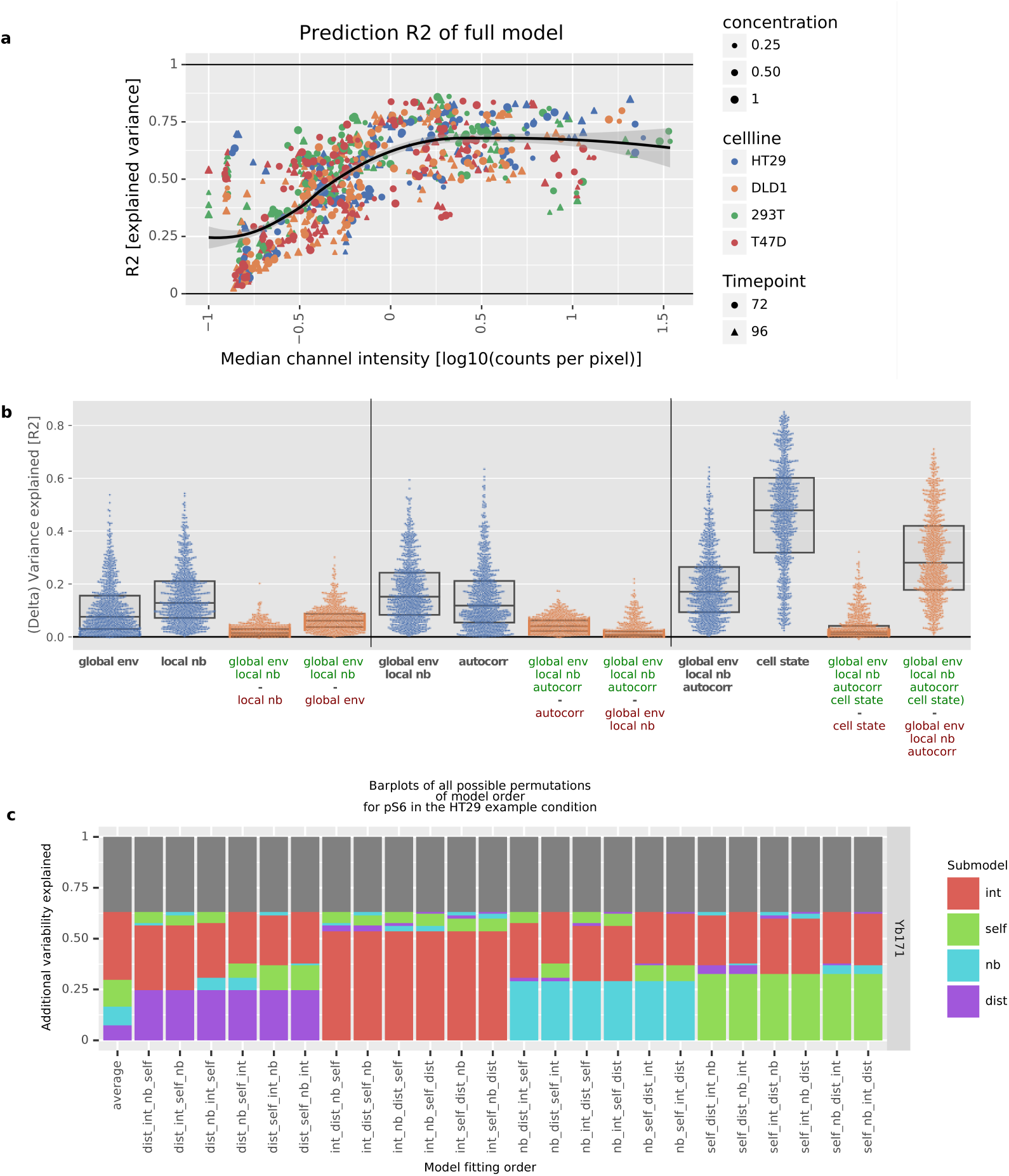
**a** Variance of each marker explained by the full linear model plotted versus median marker level. Data are shown for all cell lines, growth conditions, and growth times. **b** Variance of each marker explained by various submodels (blue dots, legend: model elements) or difference of variance explained between submodels (orange dots, legend: subtraction of variability of model with red modules from variability of model with green modules). Each data point represents a single marker from one of the 24 growth conditions (4 cell lines, 2 timepoints, 3 concentrations). **c** Barplots as in Fig. 4 but calculated from all possible orders of adding the submodules. The order is indicated in the x axis label. The bar marked ‘average’ is the contributions averaged over all model orders. Modules: dist: global environment, nb: local neighborhood, self: autocorrelation, int: cell state. Example data shown for the marker pS6 of the HT29 growth condition used for Fig. 2c (96 h growth, largest size).

**Figure S5.**
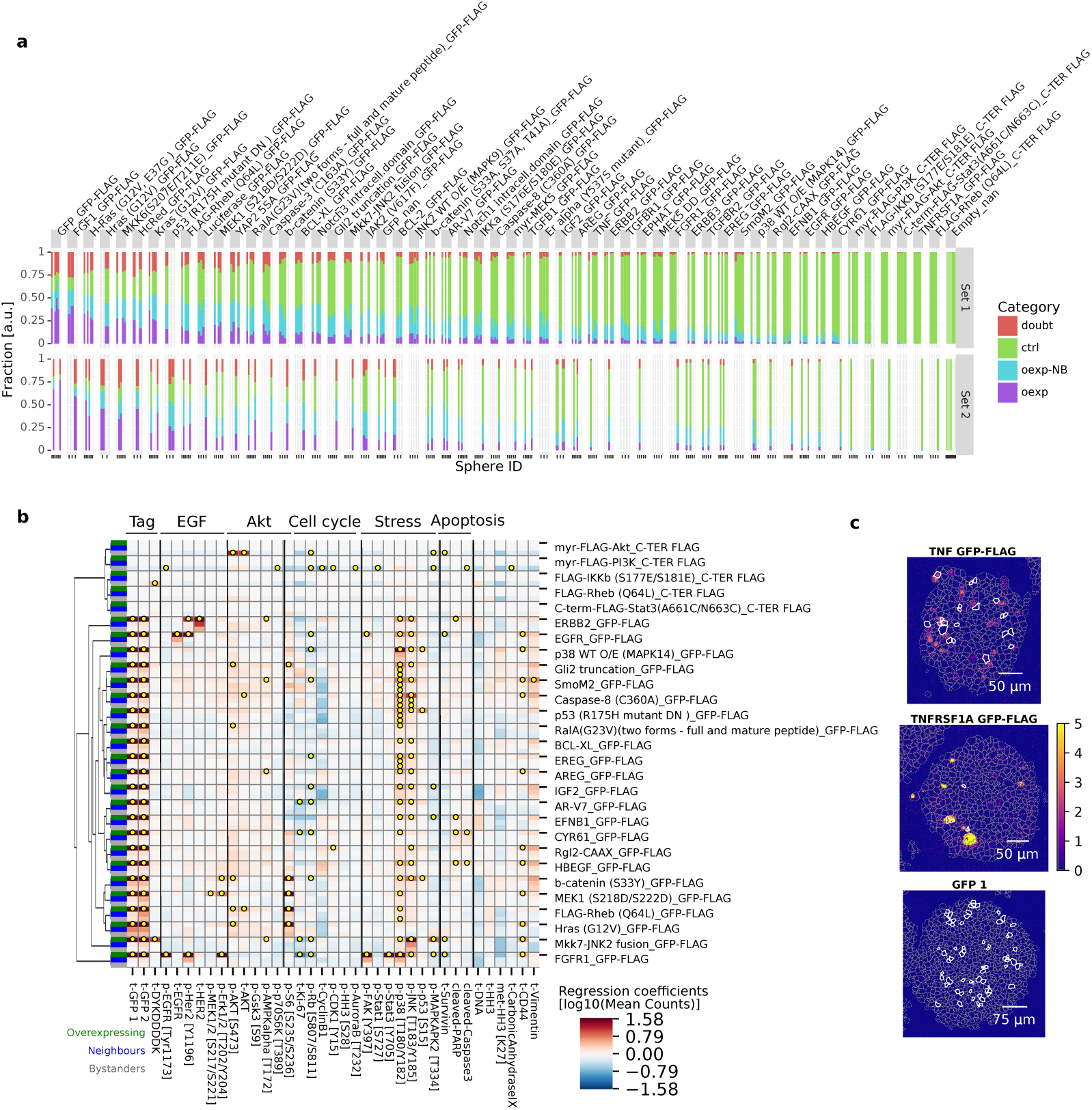
**a** Fraction of overexpressing (violet), neighbors (blue), bystanders (green), or not assigned cells (red) for each sphere replicate (bar) per overexpression construct (facet). Constructs were sorted by average fraction of overexpressing cells. **b** Matrix of overexpression constructs with intracellular effects only (rows) versus all effects on markers (columns) in cells classified as intracellular (blue), neighborhood (green), and bystander (gray) (indicated in column at the far left). Yellow dots indicate significant effects (p<0.01, q<0.1, FC > 20%). **b** Example IMC images of effects on cleaved PARP in spheroids with cells that overexpress TNF, TNFRSF1A, or GFP control. Color indicates counts per pixel. Grey dashed lines indicate segmentation mask borders and white lines mark cells identified as overexpressing.

**Table S3.**
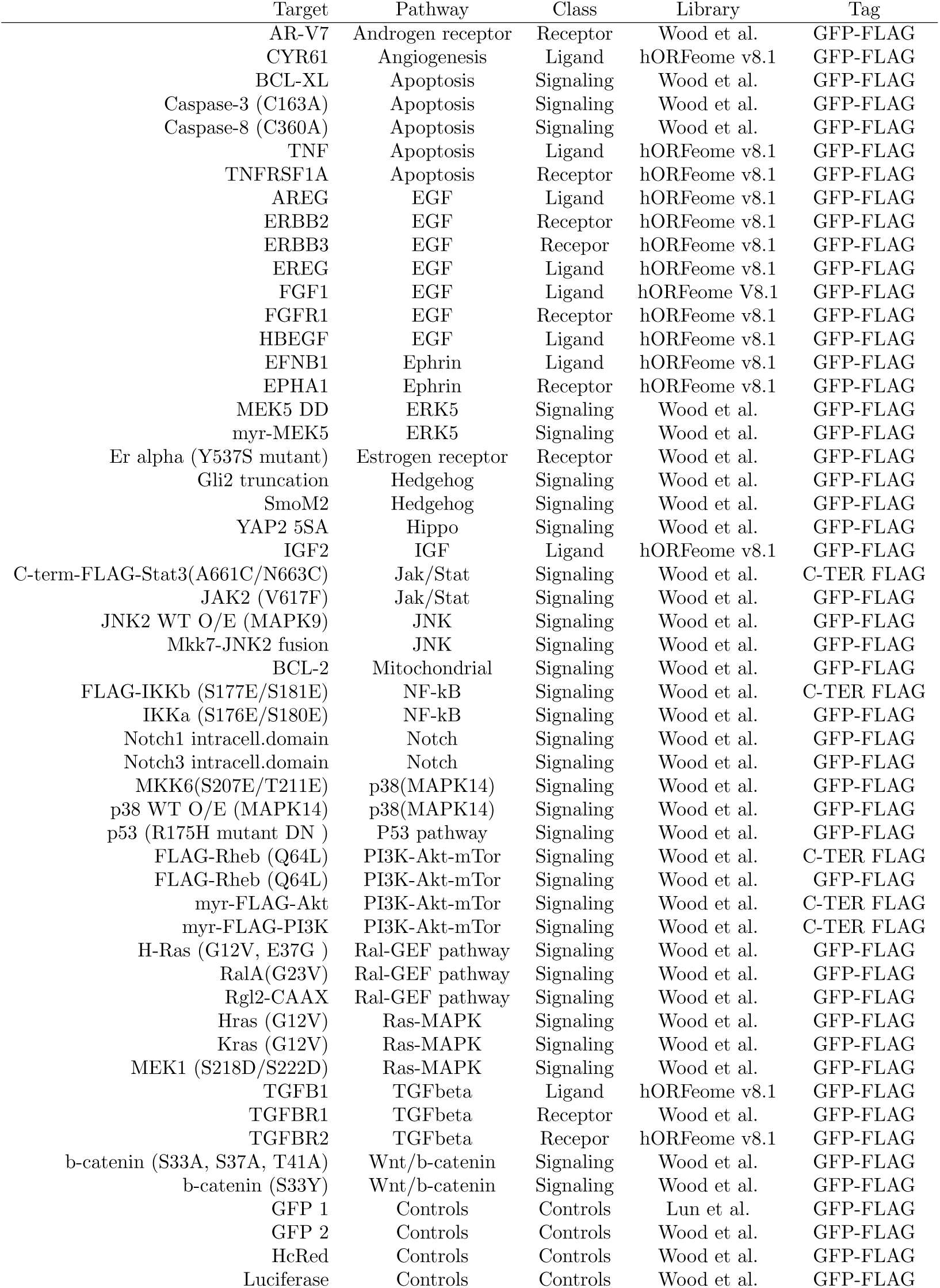
Overexpression constructs used.

